# Analyses of Biomarker Traits in Diverse UK Biobank Participants Identify Associations Missed by European-centric Analysis Strategies

**DOI:** 10.1101/2020.09.02.279844

**Authors:** Quan Sun, Misa Graff, Bryce Rowland, Jia Wen, Le Huang, Moa P. Lee, Christy L. Avery, Nora Franceschini, Kari E. North, Yun Li, Laura M. Raffield

## Abstract

Despite the dramatic underrepresentation of non-European populations in human genetics studies, researchers continue to exclude participants of non-European ancestry, even when these data are available. This practice perpetuates existing research disparities and can lead to important and large effect size associations being missed. Here, we conducted genome-wide association studies (GWAS) of 31 serum and urine biomarker quantitative traits in African (n=9354), East Asian (n=2559) and South Asian (n=9823) UK Biobank participants ancestry. We adjusted for all known GWAS catalog variants for each trait, as well as novel signals identified in European ancestry UK Biobank participants alone. We identify 12 novel signals in African ancestry and 3 novel signals in South Asian participants (p<1.61 × 10^−10^). Many of these signals are highly plausible and rare in Europeans (1% or lower minor allele frequency), including *cis* pQTLs for the genes encoding serum biomarkers like gamma-glutamyl transferase and apolipoprotein A, *PIEZ01* and *G6PD* variants with impacts on HbA1c through likely erythocytic mechanisms, and a coding variant in *GPLD1*, a gene which cleaves GPI-anchors, associated with normally GPI-anchored protein alkaline phosphatase in serum. This work illustrates the importance of using the genetic data we already have in diverse populations, with many novel discoveries possible in even modest sample sizes.

## Introduction

Lack of representation of diverse global populations is a major problem in human genetics research. As recently reviewed, 78% of genome-wide association study (GWAS) participants are of European ancestry, with an additional 9% East Asian participants[1]. All other populations (as well as multi-ethnic studies) make up less than 13% of subjects, but account for 38% of significant associations in the GWAS catalog, demonstrating the scientific importance of inclusion of diverse populations for understanding the biology of complex traits. For example, only 2.4% of GWAS participants are of predominantly African ancestry, but 7% of GWAS catalog associations were found in these participants. Inclusion of diverse populations is also essential for risk prediction; polygenic risk score instruments often perform poorly when trained using European only summary statistics and then applied to non-European populations.[2] As polygenic risk prediction moves into clinical use, this lack of representation risks perpetuating existing health disparities. Lack of inclusion of diverse populations could also lead to us missing many of the important insights into disease biology possible through human genetics.

However, as recently reviewed [3], we are failing to even use the data we have in ancestrally diverse populations. Even when data are available, many researchers focus only on large European sample sizes, and do not perform trans-ethnic or stratified analyses in those with non-European genetic ancestry. For example, the UK Biobank data, which is widely used due to its large sample size, broad data availability for qualified researchers, and variety of measured phenotypes and electronic health record data, includes >20,000 participants with non-European genetic ancestry. However, all 29 of the first papers indexed on the GWAS catalog that include UK Biobank participants included only the European ancestry sample (>400,000 individuals), likely for reasons of analytical convenience. Publicly posted summary statistics for many UK Biobank phenotypes, including serum and urine biomarkers, have recently been made available in non-European ancestry UK Biobank participants on the Pan-UK Biobank website, which provides a valuable resource for researchers [4]. However, little detailed examination of phenotypes in non-European ancestry UK Biobank participants has currently been performed.

Existing studies support the value of including even small numbers of non-European ancestry participants, especially for biomarkers and endophenotypes for which a larger percentage of variance is often explained by a small number of genetic signals. Notably, in recent trans-ethnic analyses of blood cell traits including the UK Biobank data and other cohorts (total N= 746,667), an *IL7* coding variant associated with lymphocyte counts was identified in South Asian UK Biobank participants only (N= 8189)[5]. The lymphocyte increasing allele of this variant was found to increase secretion of IL7 by 83% in follow-up *in vitro* analyses. We here assess the genetic contributors to the UK Biobank serum biomarker panel, chosen as model quantitative traits with a higher probability of previously undetected large effect size loci versus dichotomous disease endpoints. Initial analyses of these serum biomarkers have, similar to many other analyses in the UK Biobank, focused on European ancestry individuals alone[6]. This work has revealed important relationships, such as improved prediction of disease in the independent FinnGenn cohort for multi-biomarker polygenic risk scores versus single-disease PRS, particularly for liver and renal disease, and novel signals, for example a number of low frequency coding variants with impacts on kidney biomarkers and outcomes. However, we hypothesized that important novel variants were missed by the focus on European ancestry samples alone. Mendelian randomization analyses suggest causal roles for a number of these biomarkers, including IGF-1[7], urine albumin[8], urate[9], so such ancestry specific variants may have important health consequences, as well as point to key genes and biological mechanisms relevant across populations.

## Results

All genome-wide significant variants are displayed in Table 1, Figure S2 (LocusZoom[10] plots), and Table S5. We assessed at each locus whether any genome-wide significant signals remained after adjusting for the sentinel variant in Table S5. We identify 3 novel findings in South Asians (n=9823), two findings for HbA1c (a non-coding variant near *PIEZ01* and a *G6PD* missense variant) and a noncoding variant near *LPAL2* associated with Lp(a). We identify 12 novel findings in African ancestry individuals (N=9354), again including a number of coding variants (for example, *FUT6*, *GPLD1*, and *CD36* coding variants with ALP) and cis pQTLs (rare African specific noncoding variant rs541102880 with APOA, at the *APOA* gene cluster, rs57719575 at *GGT1* for liver enzyme gamma-glutamyl transferase (GGT), rs201082887 at *GPT*, also known as ALT, for alanine aminotransferase). We do not identify any novel findings in East Asians (N=2559), the smallest of the three samples. As shown in Table S5, all novel variants are rare or low frequency in Europeans. After conditioning on the sentinel variants in Table S5, additional signals were identified for several traits: at the *GPLD1* locus with alkaline phosphatase in African ancestry participants and at *LPA* in African ancestry participants and South Asian ancestry participants (Table S6, Figure S3).

**Table 1:** Novel Association Signals in African and South Asian Ancestry Participants in UK Biobank. All biomarkers are measured in serum, except creatinine, potassium, and sodium, which were measured in urine. EAF, effect allele frequency, APOA, apolipoprotein A, APOB, apolipoprotein B, ALP, alkaline phosphatase, ALT, alanine aminotransferase, BRB, bilirubin, CysC, cystatin C, GGT, gamma glutamyltransferase, HbA1c, glycated hemoglobin, IGF-1, Insulin-like growth factor 1, LPA, Lipoprotein-A

| rsID | Effect allele | Trait | Cohort | EAF | Unconditioned Results |  | Conditional Analysis |  | Nearest Gene | Annotation |
| --- | --- | --- | --- | --- | --- | --- | --- | --- | --- | --- |
| | | | | | p-value | $\beta$ | p-value | $\beta$ | | |
| rs541102880 | A | APOA | AFR | 0.1% | 2.88E-11 | 1.56 | 1.36E-11 | 1.64 | *APOA1/4/5* | noncoding |
| rs28362286 | A | APOB | AFR | 0.9% | 3.14E-20 | -0.75 | 1.28E-20 | -0.80 | *PCSK9* | coding,<br>p.Cys679Ter |
| rs146351134 | A | ALP | AFR | 0.3% | 4.64E-12 | -0.91 | 8.66E-13 | -0.95 | *GPLD1* | coding,<br>p.Trp182Cys |
| rs3211938 | G | ALP | AFR | 10.3% | 9.80E-15 | -0.20 | 8.00E-15 | -0.20 | *CD36* | coding,<br>p.Tyr325Ter |
| rs17855739 | T | ALP | AFR | 30.1% | 1.67E-19 | 0.15 | 5.87E-11 | 0.18 | *FUT6* | coding,<br>p.Glu247Lys |
| rs201082887 | G | ALT | AFR | 9.4% | 3.17E-25 | -0.27 | 1.37E-14 | -0.33 | *GPT* | noncoding |
| rs1050828 | T | BRB total | AFR female | 14.4% | 4.19E-38 | 0.31 | 1.61E-33 | 0.30 | *G6PD* | coding,<br>p.Val98Met |
|  |  |  | AFR male | 7.6% | 3.91E-33 | 0.65 | 8.85E-32 | 0.66 |  |  |
|  |  |  | AFR meta | 11.5% | 3.16E-63 | 0.36 | 8.52E-57 | 0.36 |  |  |
|  |  | BRB direct | AFR female | 14.4% | 8.04E-19 | 0.24 | 4.13E-15 | 0.21 |  |  |
|  |  |  | AFR male | 7.6% | 4.86E-20 | 0.50 | 3.31E-20 | 0.53 |  |  |
|  |  |  | AFR meta | 11.5% | 5.76E-33 | 0.38 | 2.28E-28 | 0.27 |  |  |
| rs334 | A | Creatinine | AFR | 6.3% | 2.62E-38 | -0.43 | 2.62E-38 | -0.43 | *HBB* | coding,<br>p.Glu7Ala |
|  |  | Potassium | AFR |  | 2.84E-32 | -0.39 | 2.84E-32 | -0.39 |  |  |
|  |  | Sodium | AFR |  | 5.43E-36 | -0.42 | 5.43E-36 | -0.42 |  |  |
| rs112902560 | T | CysC | AFR | 4.7% | 1.92E-11 | 0.25 | 1.92E-11 | 0.25 | *MMP26/HBB* | noncoding |
| rs57719575 | C | GGT | AFR | 14.9% | 3.97E-38 | -0.28 | 5.81E-25 | -0.25 | *GGT1* | noncoding |
| rs556126054 | G | HbA1c | SAS | 2.4% | 1.02E-27 | -0.59 | 2.19E-25 | -0.59 | *PIEZO1* | noncoding |
| rs5030868 | A | HbA1c | SAS female | 1.7% | 6.98E-22 | -0.82 | 1.56E-21 | -0.82 | *G6PD* | coding,<br>p.Ser218Phe |
|  |  |  | SAS male | 0.7% | 1.09E-33 | -1.77 | 8.79E-34 | -1.77 |  |  |
|  |  |  | SAS meta | 1.2% | 7.51E-48 | -1.06 | 1.9E-47 | -1.06 |  |  |
| rs34680334 | A | IGF-1 | AFR | 24.8% | 2.65E-15 | -0.14 | 2.24E-17 | -0.16 | *IGFALS* | coding,<br>p.His27Tyr |
| rs41270996 | T | LPA | AFR | 1.2% | 5.78E-18 | 0.66 | 1.52E-19 | 0.77 | *LPA* | noncoding |
| rs73785605 | C |  |  |  | 5.78E-18 | 0.66 |  |  |  | noncoding |
| rs374112269 | T | LPA | SAS | 0.7% | 1.83E-16 | 0.74 | 1.13E-14 | 0.68 | *LPAL2* | noncoding |

The conditional analysis p-value for our novel signals displayed is, along with GWAS catalog variants, adjusted for any variants within 1MB on each side of the sentinel variant which were genome-wide significant in analyses of serum and urine biomarkers in UK Biobank Europeans[6], to ensure the signals we identify could not be found in European ancestry participants alone.

## Discussion

Even in the relatively small number of African and South Asian ancestry individuals in UK Biobank, we identified novel and clinically relevant associations. These novel findings highlight the importance of X chromosome analysis and identify highly plausible ancestry differentiated *cis* pQTLs and coding variants.

The X chromosome is left out of the majority of GWAS analyses, with only around a third of papers including chromosome X in analysis [3, 11]. We here identify a strong association of *G6PD* coding variants with total and direct bilirubin, which has not yet been reported in the GWAS catalog despite the strong effect size. Direct bilirubin assesses bilirubin conjugated with glucuronic acid, which is secreted into bile. Indirect bilirubin (unconjugated) in plasma is usually low in healthy individuals, as this conjugation process is quite efficient, but can be elevated in many forms of hyperbilirubinemia, such as those caused by hemolysis, Gilbert syndrome, or in response to some medications[12]. This G6PD signal is also associated with indirect bilirubin (calculated as total minus direct bilirubin, β=0.22, p= 1.39e-16, females, β=0. 58, p=3.57e-24 males, β=0.29, p=1.71E-32 meta-analysis), concordant with the known risk of hemolytic anemia in those with G6PD deficiency. This association is concordant with existing literature demonstrating that males with G6PD deficiency (including deficiency caused by lead *G6PD* variant rs1050828 here) are known to be at elevated risk of neonatal hyperbilirubinemia and jaundice[13], though the strong association with bilirubin in adults and in females as well as in males is less expected. Bilirubin is commonly measured in clinical settings to assess liver function or diagnose hemolytic anemia (which can occur upon exposure to triggers such as oxidative drugs or acute infections in individuals with G6PD deficiency); if used to assess liver function, it is possible that variation at *G6PD*, as well as recently reported bilirubin associations with alpha thalassemia copy number variation,[14] could interfere with accurate clinical inference. We also identify a different *G6PD* coding variant strongly associated with HbA1c in South Asians (rs5030868, 1.1% MAF in UK Biobank South Asians, noted in ClinVar for G6PD deficiency, known as the G6PD Mediterranean variant in previous literature). Unlike the *G6PD* deficiency variant common in African Americans (rs1050828, reported here for bilirubin), which has been reported to strongly influence HbA1c in this population,[15] this variant is not previously reported in the GWAS catalog. Other *G6PD* coding variants (rs76723693 in African Americans,[16] rs72554665 and rs72554664 in East Asians[17]) have also been reported to influence HbA1c. Our results are concordant with this previous literature, and add to concerns that use of HbA1c as a laboratory test in populations with a high prevalence of G6PD deficiency may lead to underdiagnosis of diabetes and poor management and prevention of complications in those with diagnosed diabetes.[18] There is some literature to suggest that G6PD-deficient patients may have an increased risk of diabetes[19] and its complications[20]; more study is needed to disentangle impacts of G6PD deficiency on diabetes diagnosis and monitoring (due to use of HbA1c) from potential impacts on disease pathogenesis, which could be influenced by inadequate monitoring of glycemic control.

Our results in many cases point to highly plausible loci and include clinically relevant variants, for example an additional novel signal for HbA1c which likely impedes accurate assessment of glycemic control in South Asians. Along with the *G6PD* variant discussed above, a conserved noncoding variant near *PIEZ01* (rs556126054, CADD score 9.72) more common in South Asian populations (4.7% in 1000G South Asians versus 0.8% in Europeans and 0.6% in admixed Americans, not found East Asian or African populations) was associated with HbA1c. *PIEZ01* encodes an erythrocyte membrane protein, and African specific variants in this protein have been associated with red blood cell dehydration and lower malaria infection risk[21]. In recent analyses of UK Biobank blood cell trait data[5], there is a strong signal in South Asians for *PIEZ01* missense variant rs563555492 (p.Leu2277Met) for higher hematocrit (p=6.09E-14), hemoglobin (p=4.69E-22), and red blood cell count (p=1.50E-11), suggesting this variant acts through an erythrocytic pathway on HbA1c. This variant is also significant in our results (p=3.63e-21, LD r^2^=0.25 in UKBB South Asians) for HbA1c. Like the *G6PD* coding variants discussed above, this noncoding signal also likely acts through erythrocytic mechanisms and will interfere with how accurately HbA1c assess glycemic control, potentially leading to disparities in diabetes diagnosis and treatment.

We identified multiple coding variants common in African ancestry individuals for the serum biomarker alkaline phosphatase (ALP), suggesting this biomarker may be particularly susceptible to ancestry specific genetic factors. First, *FUT6* coding variant rs17855739 (p.Glu274Lys), previously identified as associated with inflammatory biomarker E-selectin in African Americans[22], was identified as associated as ALP. Other variants in *FUT2*/*FUT6* locus (but not this *FUT6* coding variant) have also been associated with other serum biomarker traits, such as cancer biomarkers CEA and CA19-9[23, 24]. ALP is also used as a cancer biomarker, among other uses. We also identified an association with ALP with African ancestry specific *CD36* nonsense variant rs3211938, which has been previously associated with HDL cholesterol levels[25, 26], ECG traits[27], red cell distribution width[28], platelet count[29], and C-reactive protein[30]. This locus is under selective pressure[31], potentially from malaria, though relationships are unclear, with this nonsense variant associated with risk of cerebral malaria and higher overall malaria incidence, but lower risk of severe anemia[32]. Finally, a rare, highly conserved coding variant at glycosylphosphatidylinositol specific phospholipase D1 (*GPLD1*) (rs146351134,p.Trp182Cys) was also associated with ALP in African ancestry individuals. This variant is quite rare across global populations, but is found at 0.4% in gnomAD v2.1.1 participants of African ancestry (with no copies found in European ancestry participants). Few genetic associations with *GPLD1* are known, but this protein hydrolyzes the inositol phosphate linkage in proteins (such as many blood cell surface proteins) anchored by phosphatidylinositol glycans (GPI-anchor). ALP is one of many important GPI-anchored proteins[33], and further study of this coding variant’s effects on other GPI-anchored proteins is warranted.

Finally, our results include a number of *cis* pQTL signals, or pQTLs in known key genes for our serum biomarkers. For example, we identified a coding variant in Insulin Like Growth Factor Binding Protein Acid Labile Subunit (*IGFLAS*), which seems likely to be a true positive as it is an African specific variant impacting a key IGF binding protein, with noncoding signals already identified in analyses in Europeans from UK Biobank identified noncoding signals at the *IGFALS* locus.[34] We also further extend the literature linking sickle cell trait (or rs334) to kidney function[35, 36], including albumin to creatinine ratio in urine, with strong associations observed for urine potassium, sodium, and creatinine. These associations are robust to adjustment for hemoglobin and estimated glomerular filtration rate (eGFR). A noncoding variant (rs112902560) in LD with rs334 was also newly identified as associated with cystatin C, another kidney function measure. We identified an association of an African ancestry specific *PCSK9* stop variant already known to be associated with LDL and total cholesterol[14, 25] with apolipoprotein B, an unsurprising extension of the existing literature. Our results also include identification of multiple novel signals at the *LPA* locus for lipoprotein A, adding to the already extensive evidence of multiple distinct *cis* pQTL signals at this locus[37–39]. We were not able to adjust for KIV2-CN (copy number) in the Lp(a) region with our imputed single nucleotide variant data, which makes these distinct signals somewhat difficult to interpret. Local ancestry has also been shown to be an important covariate at the *LPA* locus in analyses of African Americans and may be a confounder of results at this locus[38]. However, in total, these highly interpretable and biologically relevant *cis* pQTL signals echo the results from recent focused analyses of urate, IGF-1, and testosterone in European populations.[34] Many lead signals for these serum biomarkers were near genes involved in biosynthesis, transport, or signaling pathways relevant to the target trait, in contrast to the often difficult to interpret lead association signals for more complex phenotypes.

Some of our novel findings would not have been possible without TOPMed imputation, which has been demonstrated in previous analyses to have dramatically improved imputation quality for rare variants, particularly in Hispanic/Latino and African ancestry individuals[40], including for identification of rare variant association signals in African[40] and European ancestry[41] UK Biobank participants. For many of our identified signals, imputation quality was similar to the Haplotype Reference Consortium (HRC) and UK10K haplotype imputation provided by UK Biobank. However, for some novel signals variants were absent from this previously used reference panel (for example *GPLD1* coding variant rs146351134) or were imputed with an info score <0.3 (*G6PD* coding variant rs5030868). We do note that due to stringent variant filtering in TOPMed some important known signals (like sickle cell trait) were not included in the reference panel; this is an important limitation for users of this reference.

Given the very large sample size now available for all of these biomarkers through unpublished analyses in European ancestry individuals in UK Biobank, as well as in many cases other large GWAS meta-analyses, it is striking that a number of functionally plausible and novel signals could be identified in analyses of <10,000 African and South Asian individuals, a sample size much smaller than most current GWAS analyses. Our results highlight the potential impact of ancestry-differentiated results on the accuracy of clinical biomarker measures. Issues with the use of HbA1c in non-European populations due to *G6PD* variants, sickle cell trait, and other ancestry differentiated variants are recognized, but other clinical assays are also likely influenced by ancestry differentiated variants unrelated to disease risk. This bias may cause even more systematic problems as novel biomarkers and large-scale proteomics panels move into clinical risk prediction, as the largest training datasets for risk prediction and determination of reference ranges are composed of European ancestry individuals. A major limitation of our results is our failure to provide replication for some of our putative novel findings, due to a lack of readily available replication datasets, especially for less frequently measured serum biomarkers (for example IGF-1, where the largest existing GWAS analysis (other than UK Biobank Europeans) includes only 10,280 European ancestry individuals[42]). However, the number of variants identified with strong functional annotation near relevant genes suggests that these preliminary results include a number of findings worthy of future exploration in larger datasets of diverse ancestry background, and clearly demonstrate the value of using genetic data from UK Biobank non-European ancestry participants.

## Materials and Methods

The UK Biobank resource includes genetic and phenotypic data on nearly 500,000 individuals aged 40-69 at time of recruitment (2006-2010).[43] The UK Biobank recently released data on 34 serum and urine biomarkers, chosen to reflect a wide range of diseases based on their role as established risk factors or diagnostic measures, with an emphasis on renal and liver health.[44] We excluded three biomarkers with a high percentage of values below the reportable range (oestradiol, microalbumin in urine, and rheumatoid factor, with missingness>70%) and assessed inverse normalized values for all other biomarker traits, leaving 31 biomarkers for genetic analysis (Table S1).

We used a combination of self-reported ancestry and k-means clustering of genetic principal components to derive lists of individuals to include in the African, South Asian, and East Asian clusters. First, we calculated principal components (PC) and their loadings for all 488,377 genotyped UKBB participants using high quality variants in the UK Biobank data set that overlapped with the participants in the 1000G Phase 3 v5 (1KG) reference panel (Figure S1). Reference ancestries used included 504 European (EUR), 347 American Admixed (AMR), 661 African (AFR), 504 East Asian (EAS), and 489 South Asian (SAS) samples (overall 2504). We projected the 1KG reference panel dataset on the calculated PC loadings from UKBB. We then used k-means clustering with 4 dimensions, defined by the first 4 PCs, to identify the individuals that clustered with the majority of 1KG reference panels in each ancestry (PC1, PC2, PC3, and PC4 are displayed in the figure below, those who are not in any kmeans cluster (UKBB_other) are shown in grey).

We used self-reported ancestry/ethnicity (variable “ethnic_background”), in some circumstances, to adjust these groups (for example to include admixed individuals who fell outside of the 1000G clusters, but still had substantial non-European ancestry). For the African ancestry subset used in our analysis, we included all individuals that cluster with the 1KG AFR samples by k-means clustering, except n=7 individuals self-reported as follows in variable Ethnic background (variable 21000-0.0), at baseline visit (due to the possibility of a sample swap): White, British, Irish, Any other White background, Indian, Pakistani, Bangladeshi, Any other Asian background, or Chinese. We also added to our African ancestry cluster those that did not cluster in a group using k-means, but self-reported White and Black Caribbean, White and Black African, Black or Black British, Caribbean, African, or Any other Black background (n=660). For the South Asian subset used in our analysis, we included all individuals that cluster with the 1KG SAS samples by k-means clustering, except n=117 individuals that self-reported as follows: White, British, Irish, Any other White background, White and Black Caribbean, White and Black African, Black or Black British, Black Caribbean, African, Any other Black background, or Chinese. We added individuals that did not cluster in a group using k-means, but self-reported Indian, Pakistani, Bangladeshi (n=55). Finally, our East Asian ancestry subset is comprised of individuals that cluster with the most 1KG East Asians (EAS) by k-means clustering, removing n=8 individuals that self-report White, British, Irish, Any other White background, White and Black Caribbean, White and Black African, Indian, Pakistani, Bangladeshi, Black or Black British, Black Caribbean, African, or Any other Black background. We also added those that fell among those that did not cluster in a group using k-means, but self-reported Chinese (n=19). After clustering and exclusion of extreme outliers/potential sample swaps, we included n=9354 African, n=2559 East Asian, and n=9823 South Asian ancestry participants. For ease of comparison to reference allele frequencies, we stratified analyses by ancestry group.

Imputation was performed using 97,256 deeply sequenced reference genomes from diverse populations from the National Heart, Lung, and Blood Institute’s Trans-Omics for Precision Medicine Initiative (https://imputation.biodatacatalyst.nhlbi.nih.gov/#!), in order to better capture ancestry-specific rare variation (particularly in African ancestry populations) versus the UK10K panel used for the public UK Biobank release. We filtered to individuals and SNPs with a call rate >90% prior to imputation. For our analyses, we assessed common variants (MAF > 0.5%) with estimated r^2^ > 0.3 and rare variants (MAF < 0.5%) with estimated r^2^ > 0.8. Association analyses were performed using EPACTS 3.3.0 using the EMMAX test, which accounts for population structure. Genotyped variants with MAF>1% and missing rate < 1% were used in kinship matrix derivation. We removed variants with an estimated minor allele count < 5 when running EPACTS to improve model stability. X chromosome analyses were conducted stratified by sex and then meta-analyzed using GWAMA, alleviating problems with inflation for some sex-differentiated biomarkers and allowing us to assess evidence of heterogeneity by sex. We assessed testosterone in a sex stratified fashion for all chromosomes due to the dramatically different distribution in males and females (see Table S1). For subjects on lipid medications, we divided total cholesterol by 0.8 to approximate pre-medication values, and we divided directly assessed LDL by 0.7, as previously recommended.[45] For analysis of both diabetes related traits, we excluded individuals with diabetes diagnosed by a doctor (field 2443-0.0), those taking insulin (field 6153-0.0), HbA1c >=48 mmol/mol, or glucose >=7 mmol/L.

For our analysis of serum and urine biomarkers, we first regressed out covariates (age, sex, 10 PCs (provided by UK Biobank), genotyping array, centers), then inverse normalized residuals. In our EPACTS models, we included known variants from the GWAS catalog (accessed Spring 2020) as covariates (any variant previously identified on each tested chromosome, Table S2), as our primary aim was to identify novel signals missed in previous predominantly European analyses. For identified signals, we evaluated if analyses in Europeans from UK Biobank alone (as described in [6] and accessed using PheWas browser at https://biobankengine.stanford.edu[46] (variants not yet available in the GWAS catalog, Table S3)) had identified genome-wide significant variants (p<5×10^−8^) within 1MB of our sentinel signal. We then included these nearby variants as covariates, if any were reported, in final conditional analyses reported here, to see whether our sentinel variants from non-European ancestry focused analyses were still genome-wide significant. Chromosome X was not included in previous European focused analyses, so this does not apply to those variants.

We did not observe evidence of significant genome-wide inflation (Table S4). We adopted a significance threshold of 5 × 10^−9^/31 traits, or p<1.61 × 10^−10^, based on reasonable estimates of the number of independent tests for testing all common and low frequency variants genome-wide[47]. Our initial analyses identified several putative novel signals at the *HBB* locus; however, these results were difficult to interpret as known sickle cell trait variant rs334, which is known to have impacts on numerous traits including kidney function[36] and HbA1c[48] was excluded from the TOPMed freeze 8 reference panel. We extracted this variant from UK10K imputation provided by UK Biobank (imputation info score 0.899) for additional conditional analyses at these loci. This variant was the lead novel signal at *HBB* after its inclusion for urine creatinine, potassium, and sodium (Table 1 and Table S5).

## Acknowledgements

This research has been conducted using the UK Biobank Resource under Application Number 25953.

LMR is supported by T32 HL129982 and KL2 TR00249. NF is supported by the NIH DK117445, MD012765 and HL140385.

We would like to acknowledge use of the Trans-Omics in Precision Medicine (TOPMed) program imputation panel (freeze 8 version) supported by the National Heart, Lung and Blood Institute (NHLBI); see www.nhlbiwgs.org. TOPMed study investigators contributed data to the reference panel, which was accessed through https://imputation.biodatacatalyst.nhlbi.nih.gov. The panel was constructed and implemented by the TOPMed Informatics Research Center at the University of Michigan (3R01HL-117626-02S1; contract HHSN268201800002I). The TOPMed Data Coordinating Center (R01HL-120393; U01HL-120393; contract HHSN268201800001I) provided additional data management, sample identity checks, and overall program coordination and support. We gratefully acknowledge the studies and participants who provided biological samples and data for TOPMed.

## Web Resources

Summary statistics are available at https://yunliweb.its.unc.edu/serum_biomarker/index.php Data are available upon request from the UK Biobank https://www.ukbiobank.ac.uk/.

