## Supplement for "Analyses of Biomarker Traits in Diverse UK Biobank Participants Identify Associations Missed by European-centric Analysis Strategies"

Table S1: Sample characteristics table, with a summary of the distribution for each quantitative measure in each population in males and females separately and both sexes combined.

Table S2: GWAS catalog variants adjusted for as covariates for each analyzed trait. Please note that for bilirubin, most previous GWAS analyses are for total bilirubin; the same set of variants is also adjusted for in analyses of direct and total bilirubin. The reported p-value is from the GWAS catalog. The minor allele frequency (MAF) and imputation R squared are listed; variants with an imputation quality <0.3 were not adjusted for and are listed as NA.

Table S3: Variants identified in analyses of European UK Biobank participants^5^ and within 1 Mb of a putative novel signal in our analyses of African, South Asian, and East Asian UK Biobank participants, and thus adjusted for in our final conditional analysis results.

Table S4: Genomic inflation lambdas for all autosomal and chromosome X analyses, adjusted for GWAS catalog and Sinnott-Armstrong et al. preprint (UK Biobank European ancestry analyses) variants. Cohorts column lists the ancestry-stratified subcohort(s) in which we adjusted for the listed variant.

Table S5: Novel association signals in African and South Asian populations. We also display the minor allele counts and frequencies in all 1000G reference populations, the CADD PHRED^48^ and FATHMM-XF^49^ scores, and the nearest gene and annotated consequence for each genome-wide significant variant. MAC, minor allele count, EAF, effect allele frequency, APOA, apolipoprotein A, APOB, apolipoprotein B, ALP, alkaline phosphatase, ALT, Alanine aminotransferase, BRB, bilirubin, CysC, cystatin C, GGT, gamma glutamyltransferase, HbA1c, glycated hemoglobin, IGF-1, Insulin-like growth factor 1, LPA, Lipoprotein-A.

Table S6: Secondary signals, conditioning on both novel sentinel variants from Supplementary Table 5 and GWAS catalog and Sinnott-Armstrong et al. preprint (UK Biobank European ancestry analyses) variants in the region. We also display the minor allele counts and frequencies in all 1000G reference populations, the CADD PHRED^48^ and FATHMM-XF^49^ scores, and the nearest gene and annotated consequence for each genome-wide significant variant. MAC, minor allele count, EAF, effect allele frequency, ALP, alkaline phosphatase, GGT, gamma glutamyltransferase, LPA, Lipoprotein-A.

Figure S1: Plots for UK biobank principal components (PC) 1-4, with UK Biobank analyzed jointly with 1000 Genomes samples for assessment of ancestry clustering using kmeans (grey samples (UKBB_other) do not fall in any kmeans cluster).

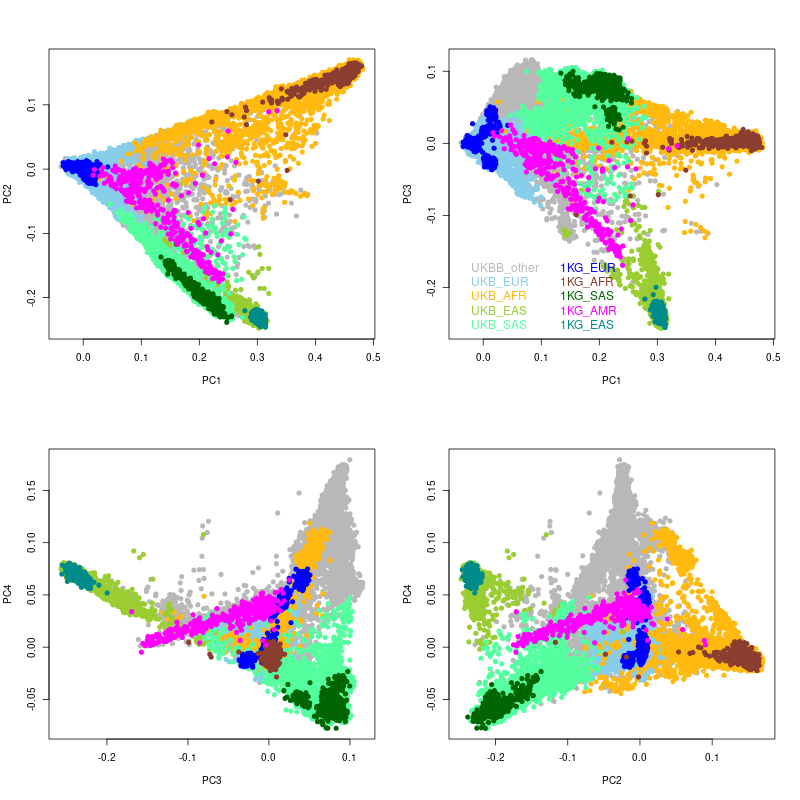

Figure S2: LocusZoom plots for all conditionally distinct signals, with all SNP association statistics conditioned on known GWAS variants for a given trait (Table S2) and Sinnot Armstrong et al. preprint (Table S3) variants within 1 Mb of novel sentinel variant. Linkage disequilibrium is calculated based on the UK Biobank samples used for a given ancestry grouping.

a. rs541102880, Apolipoprotein A (APOA), African ancestry

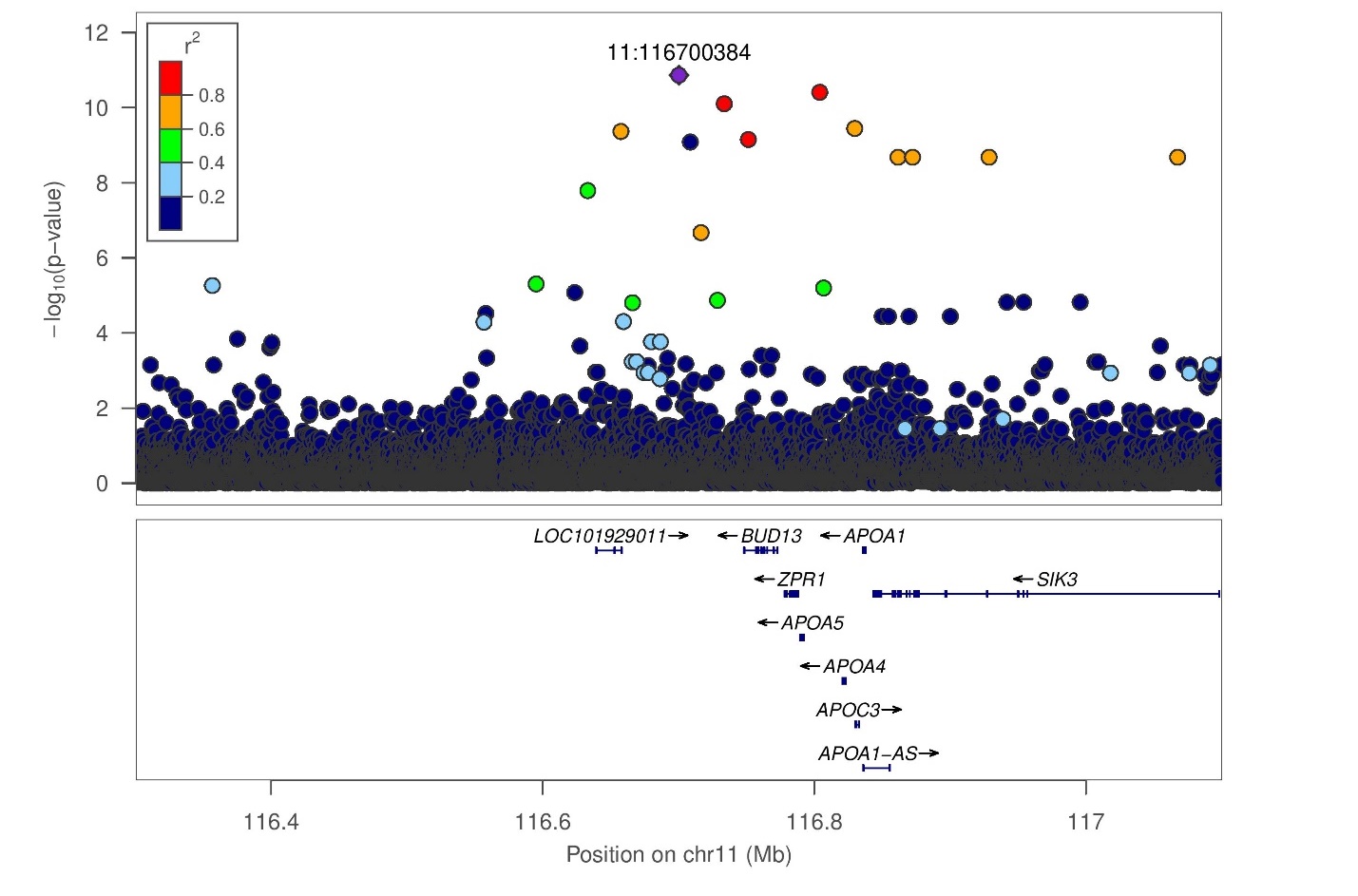

b. rs28362286, Apolipoprotein B (APOB), African ancestry

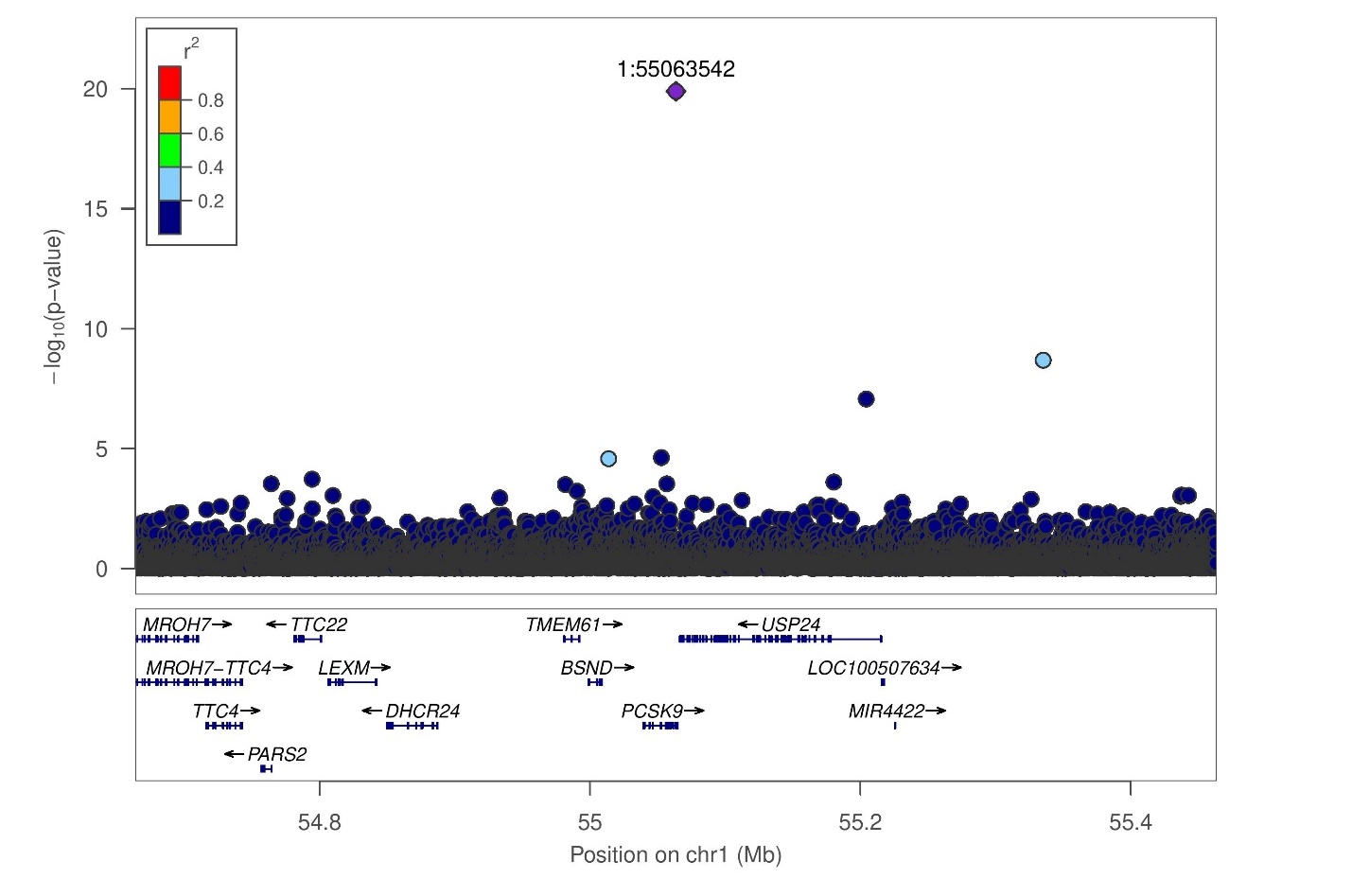

c. rs146351134, alkaline phosphatase, African ancestry

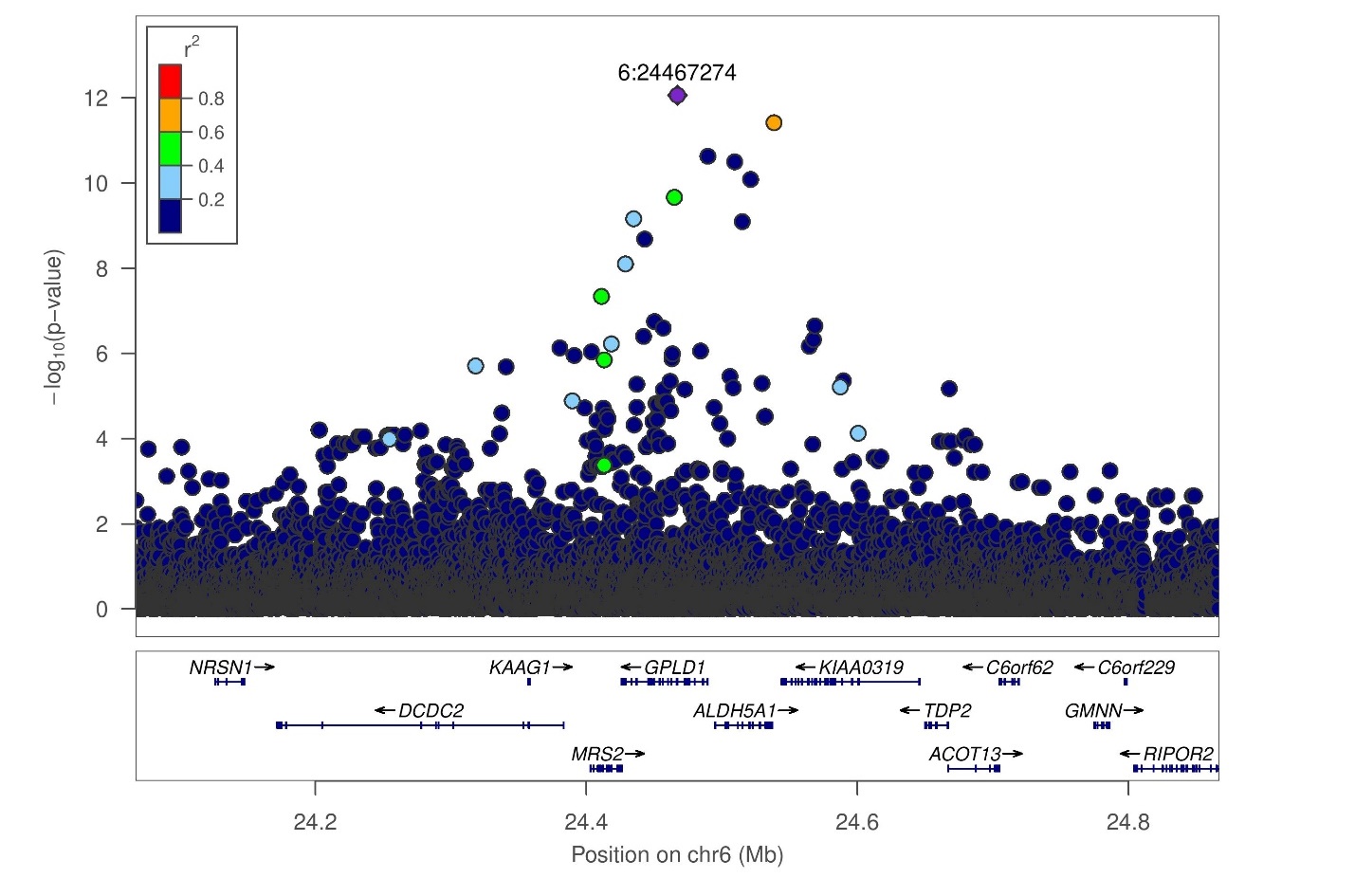

d. rs3211938, alkaline phosphatase, African ancestry

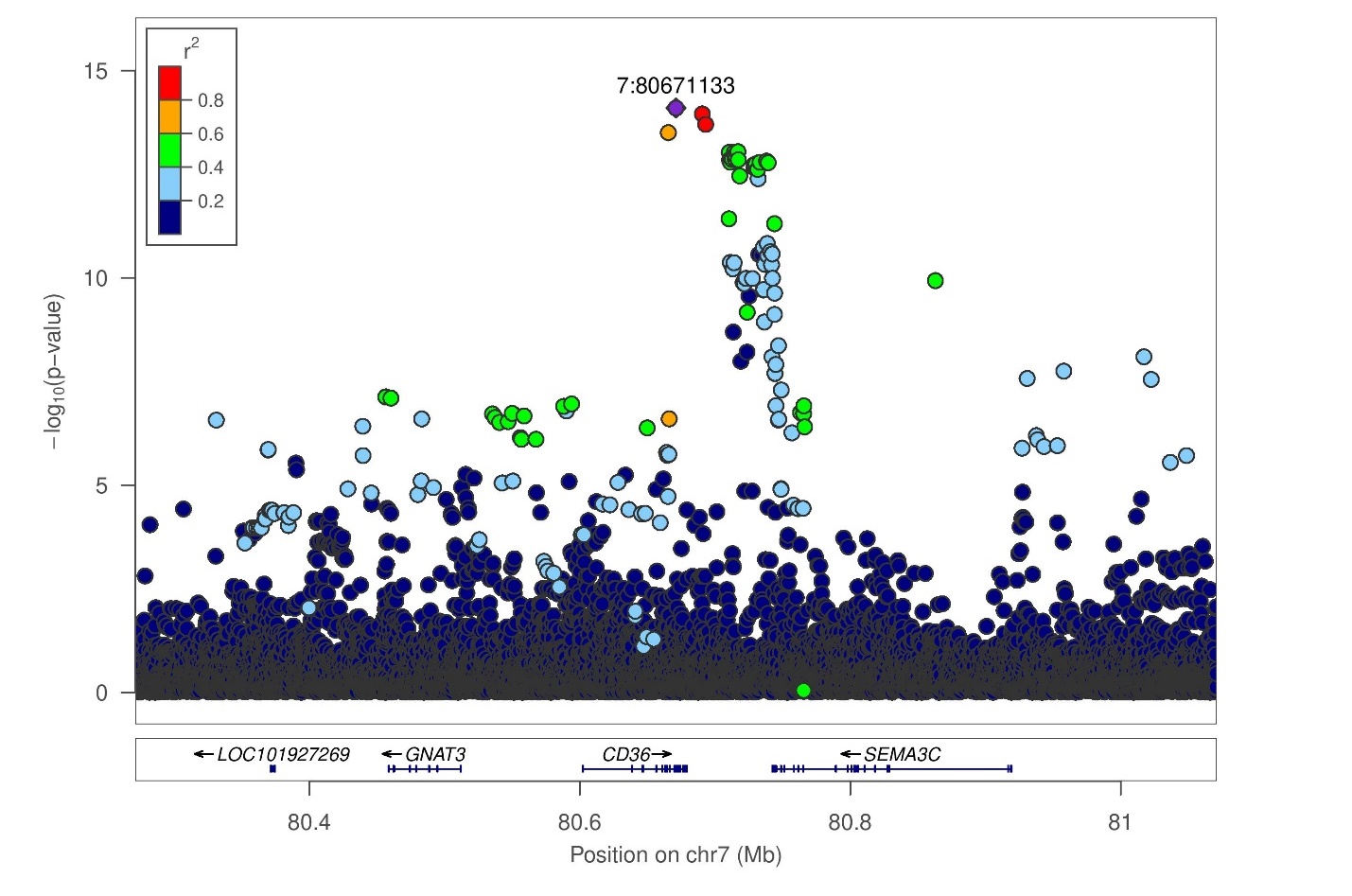

e. rs17855739, alkaline phosphatase, African ancestry

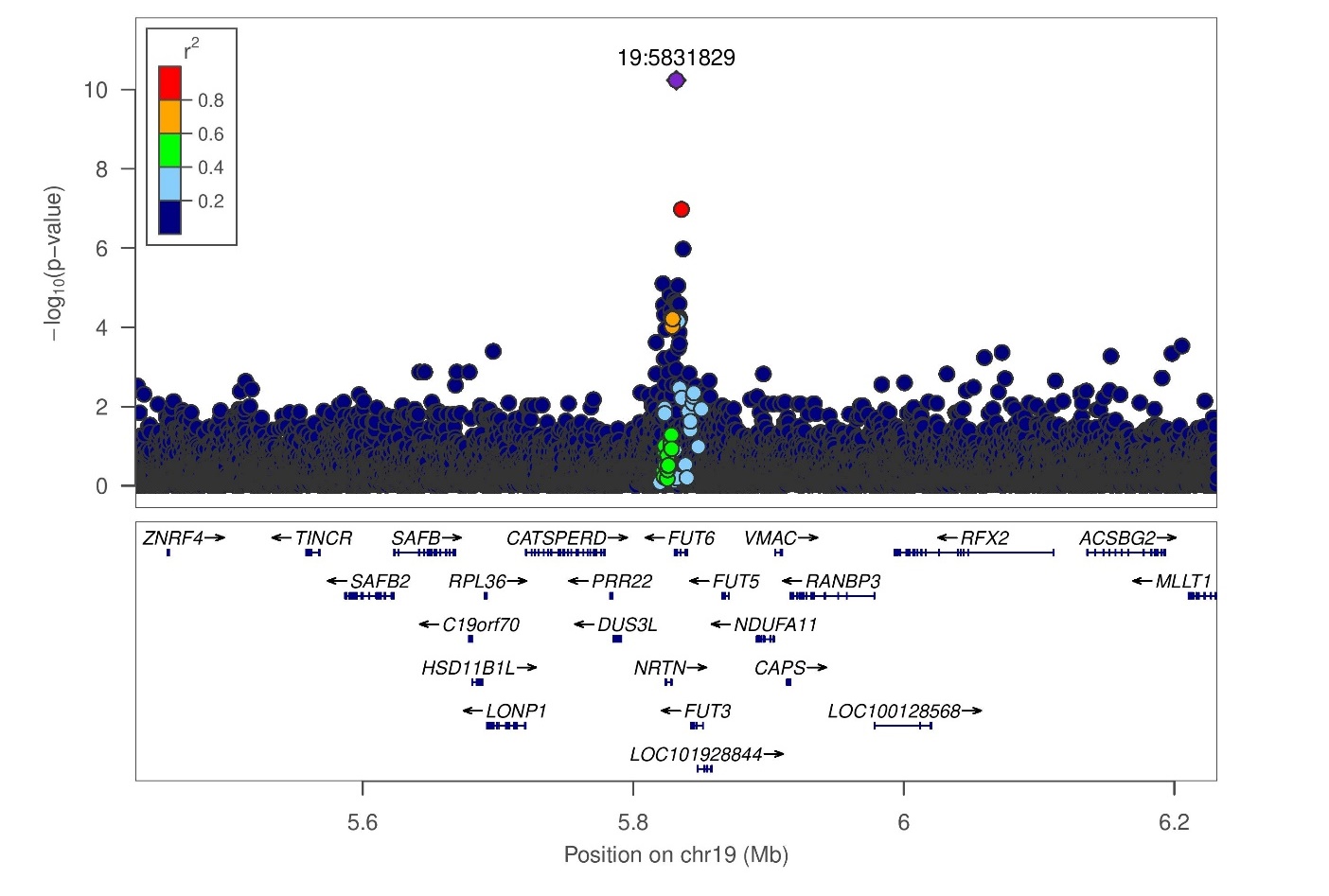

f. rs201082887, alanine aminotransferase, African ancestry

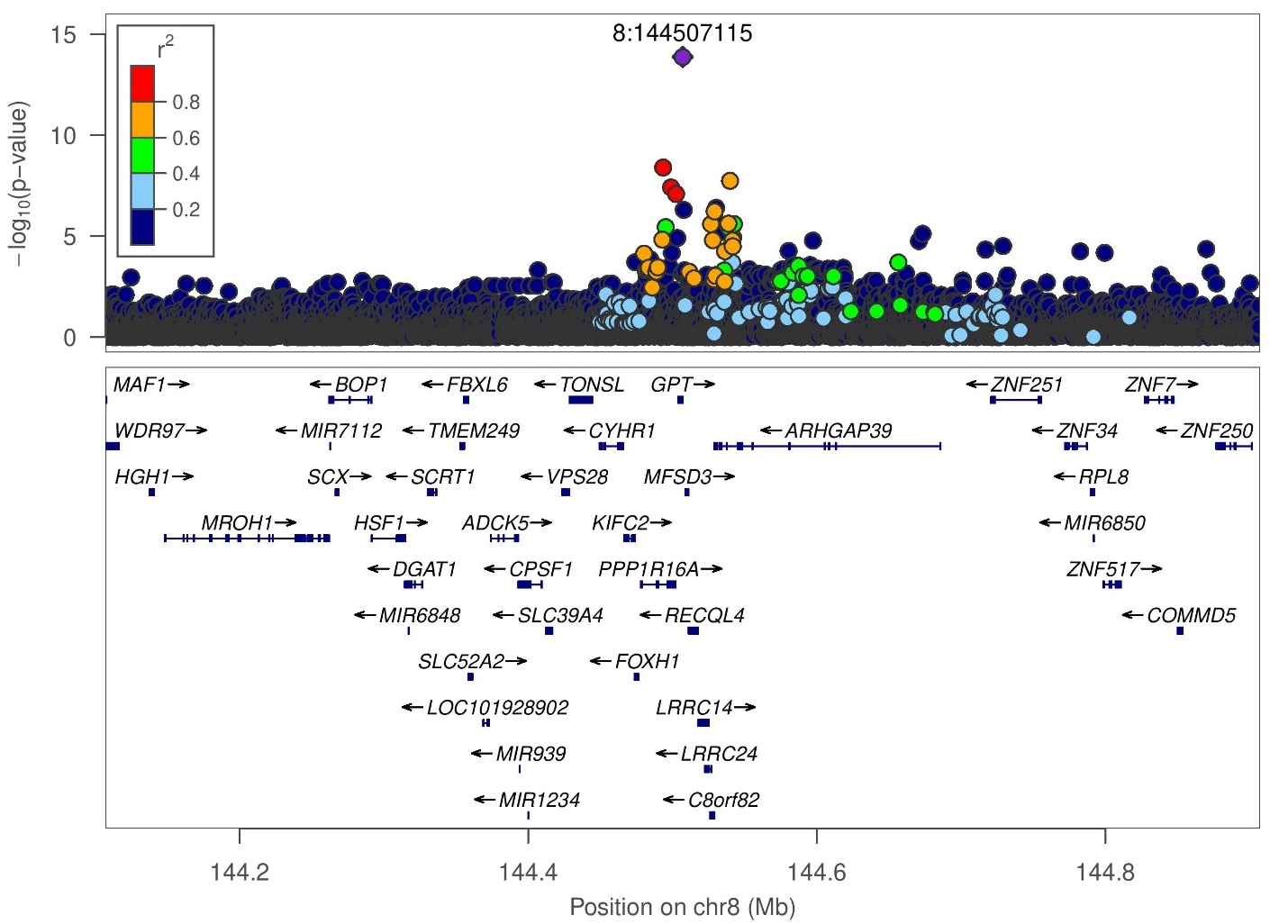

g. rs1050828, bilirubin total, African ancestry

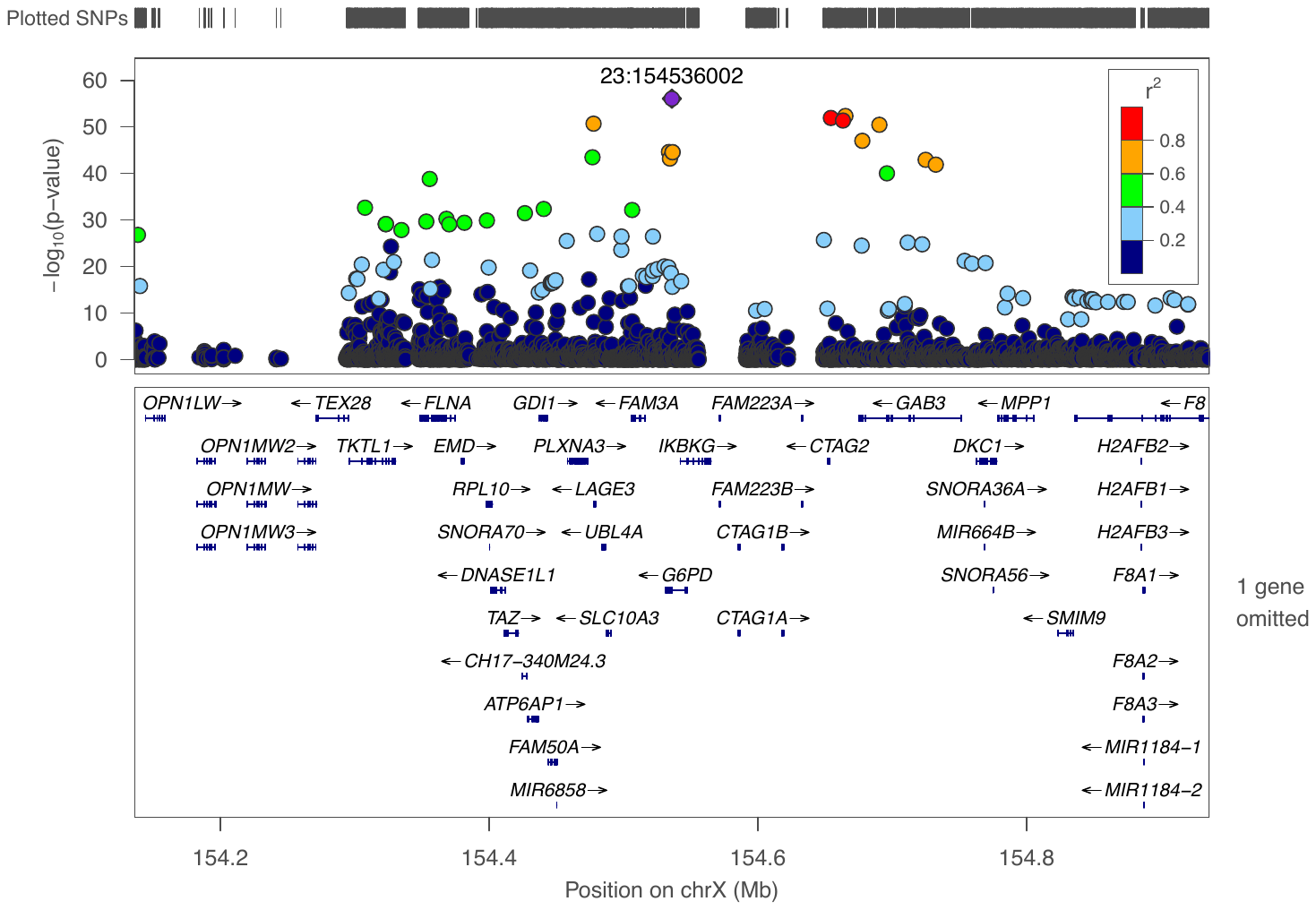

h. rs1050828, bilirubin direct, African ancestry

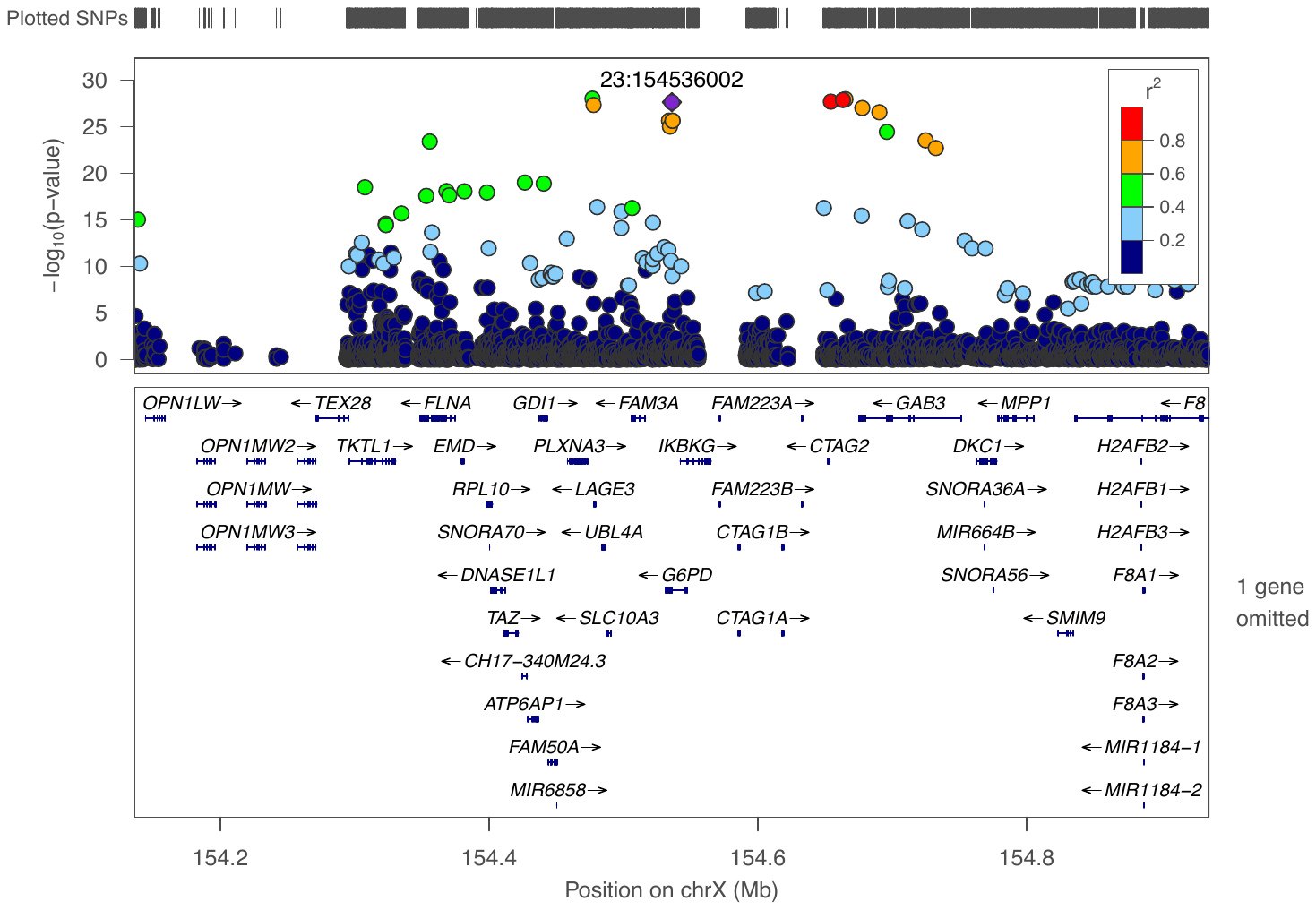

i. rs334, urine creatinine, African ancestry

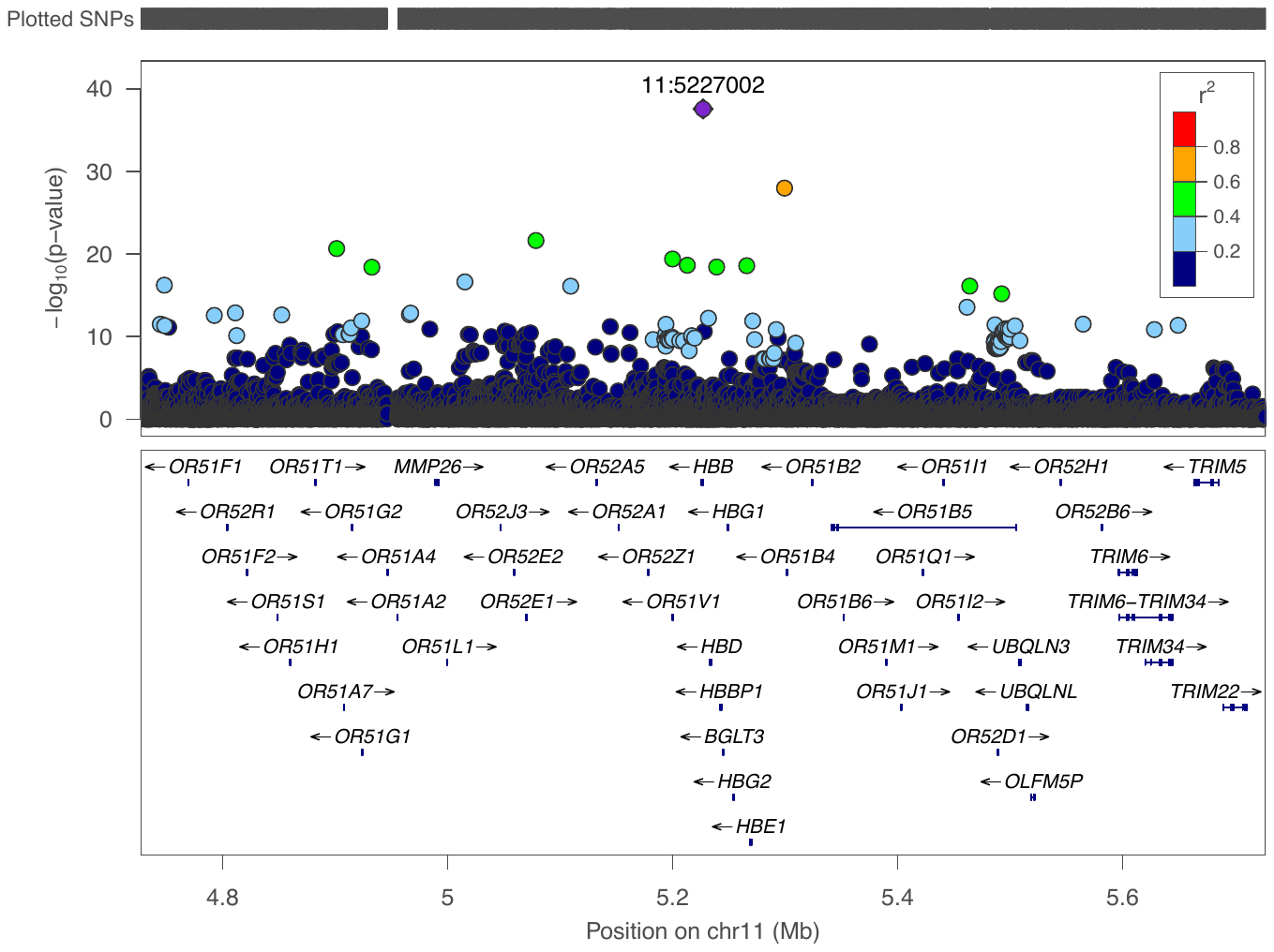

j. rs334, urine potassium, African ancestry

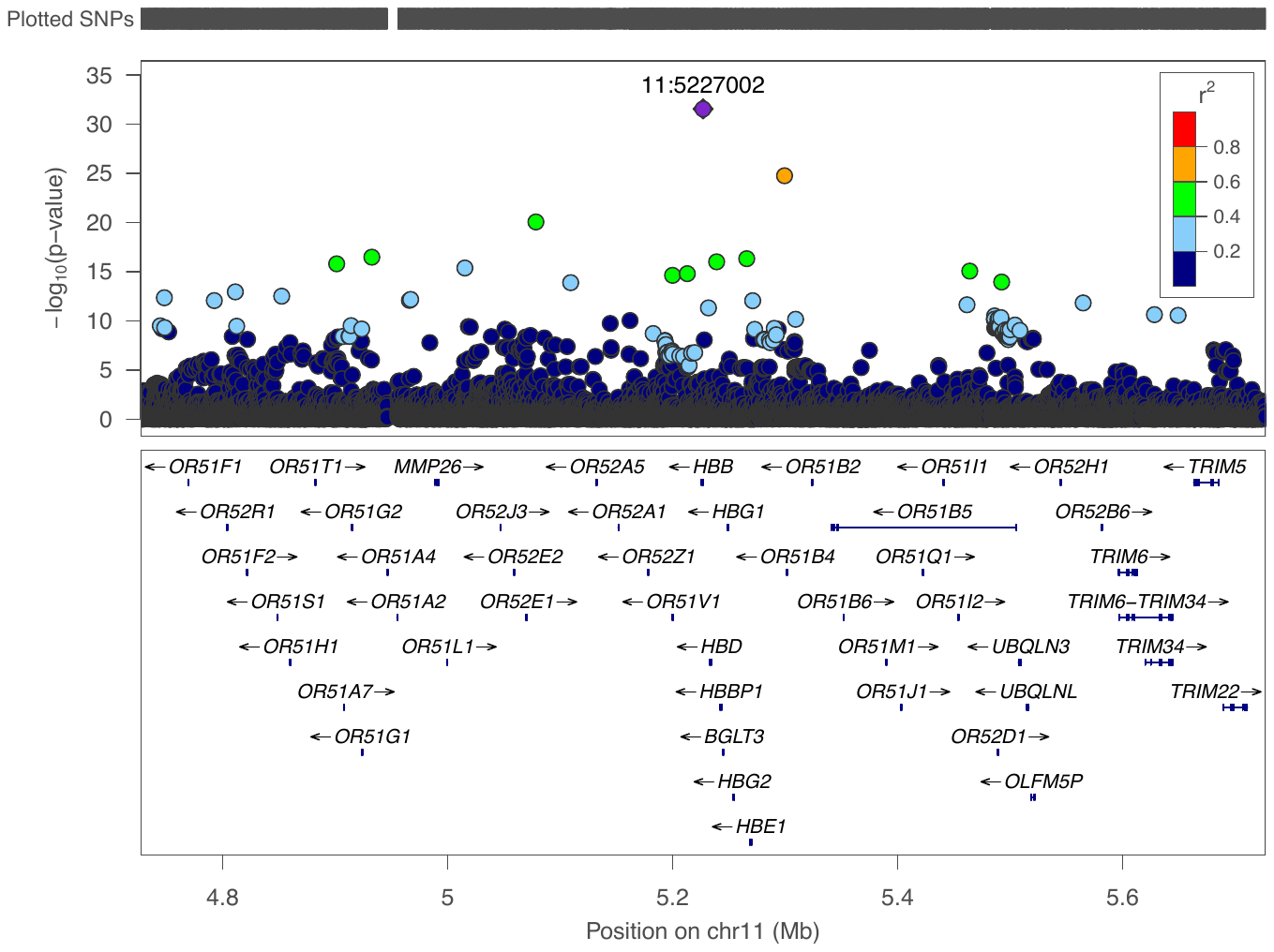

k. rs334, urine sodium, African ancestry

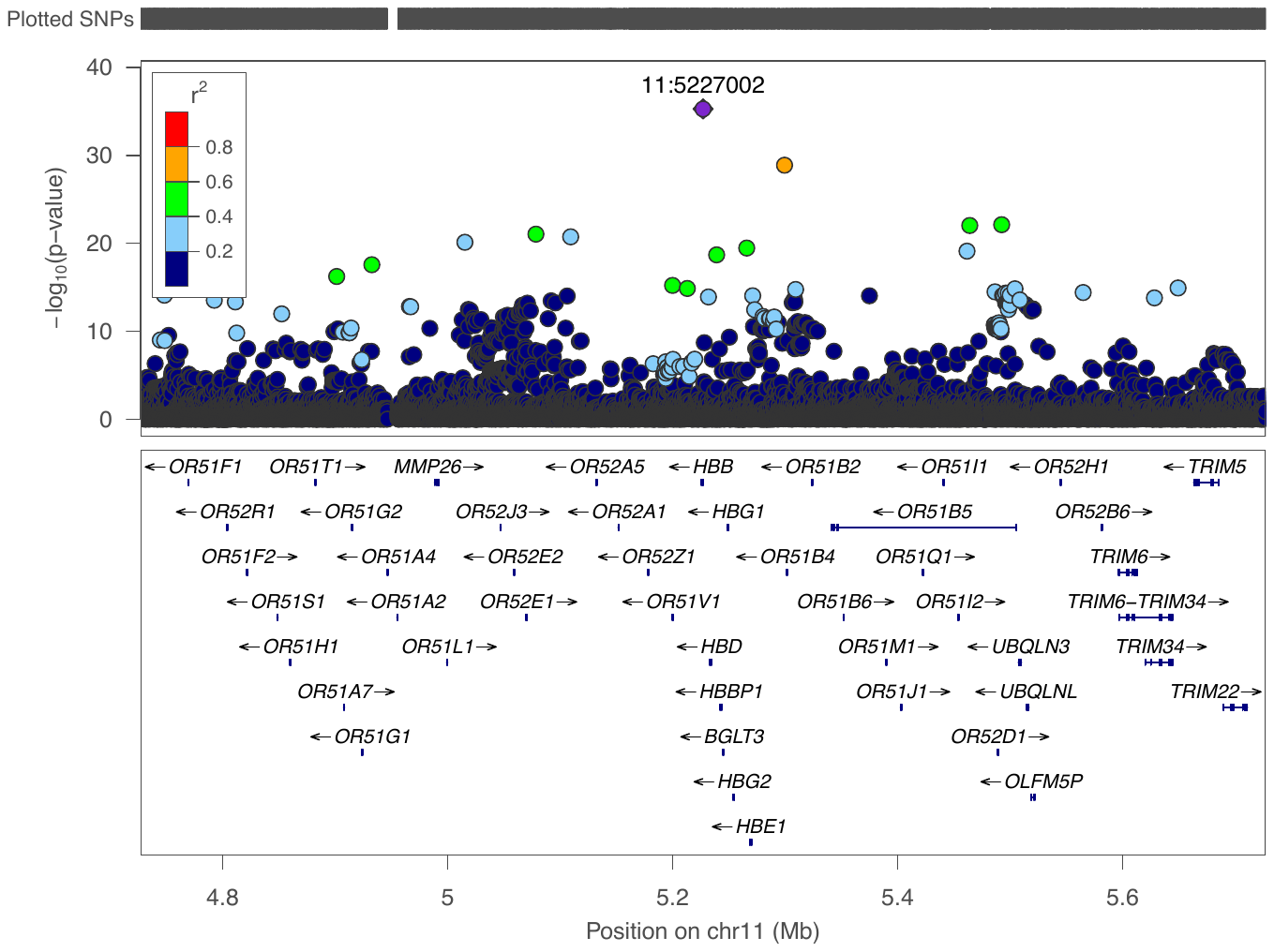

l. rs112902560, cystatin C, African ancestry

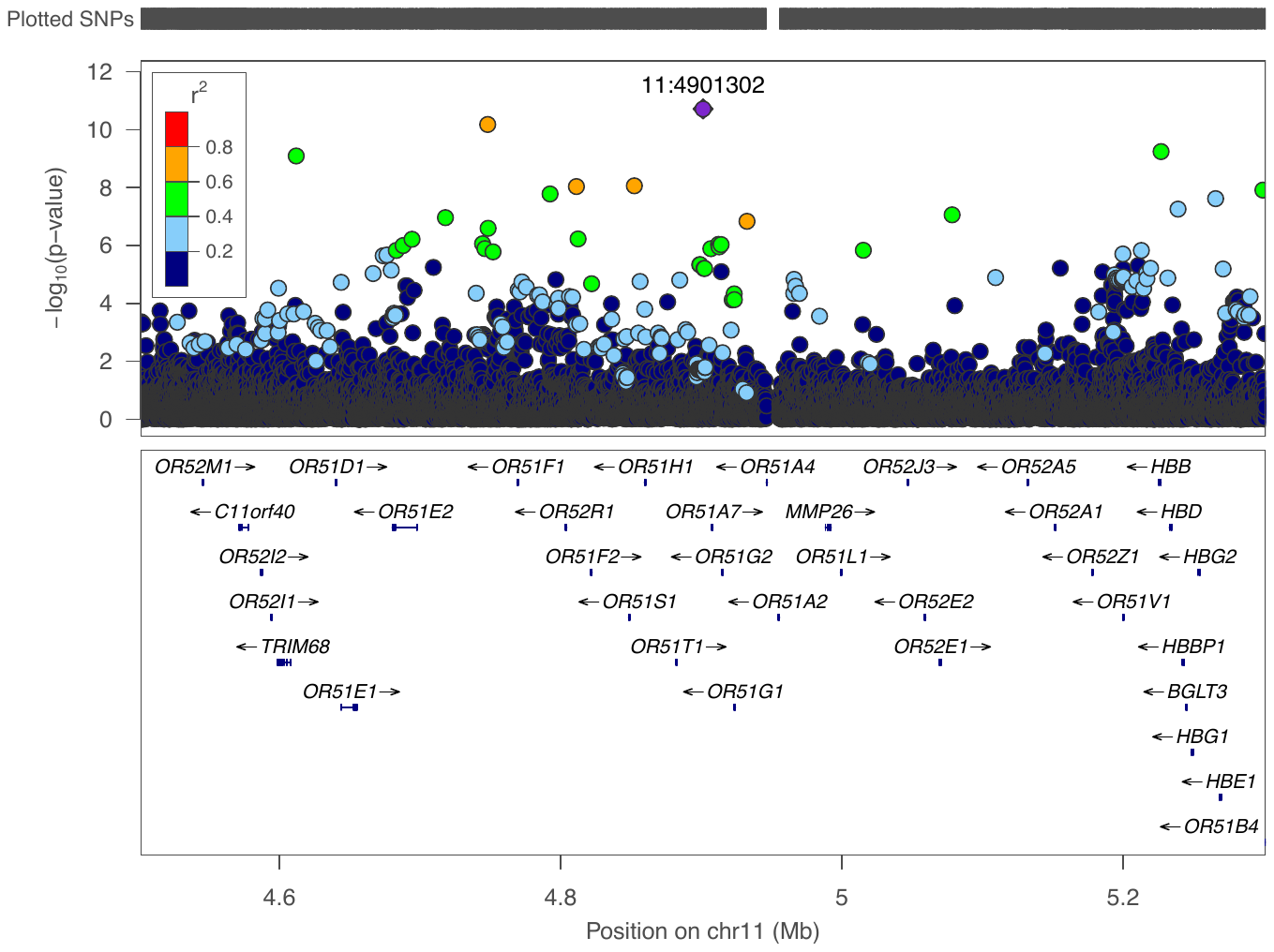

m. rs57719575, gamma glutamyltransferase, African ancestry

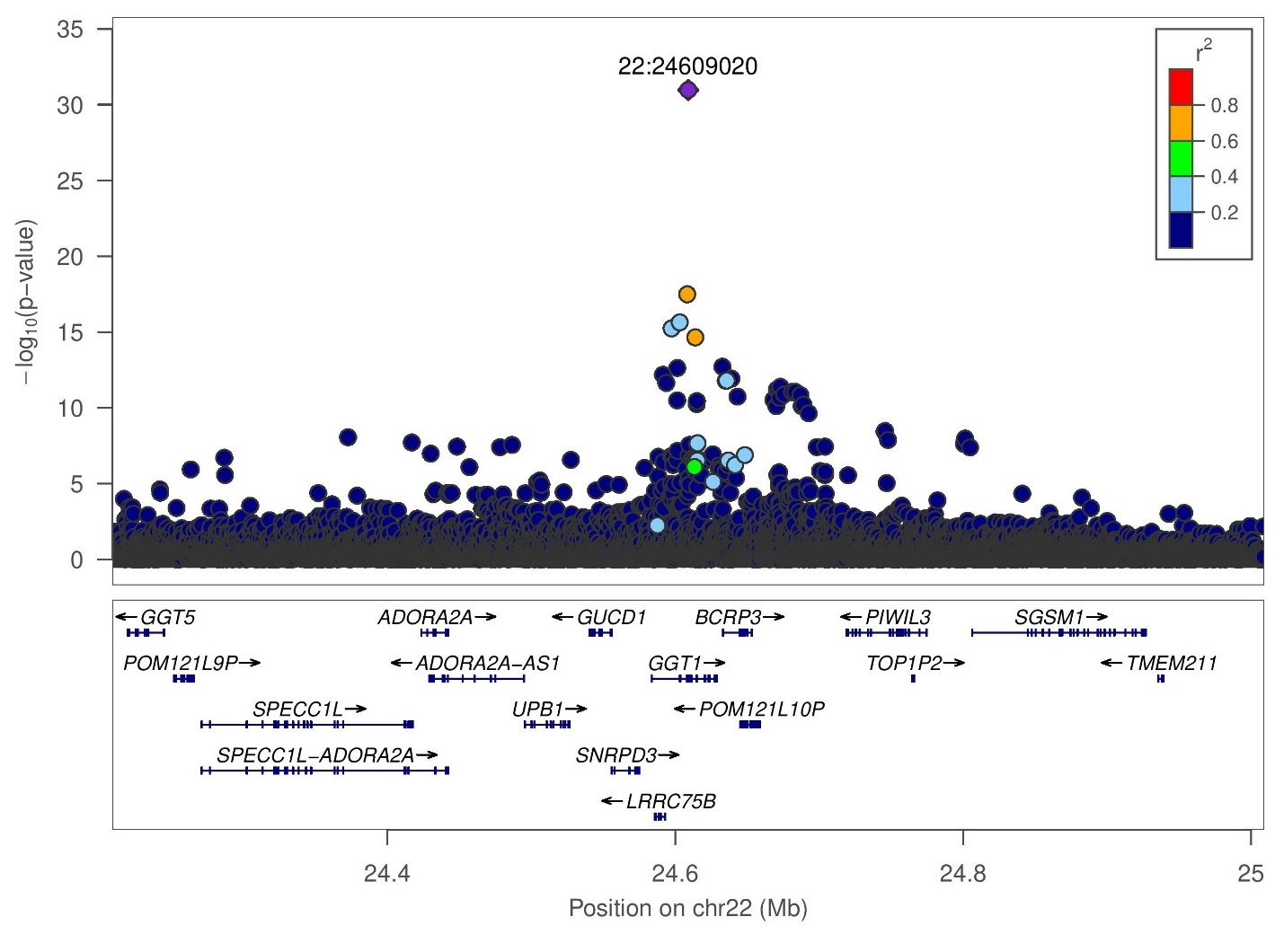

n. rs556126054, glycated hemoglobin, South Asian ancestry

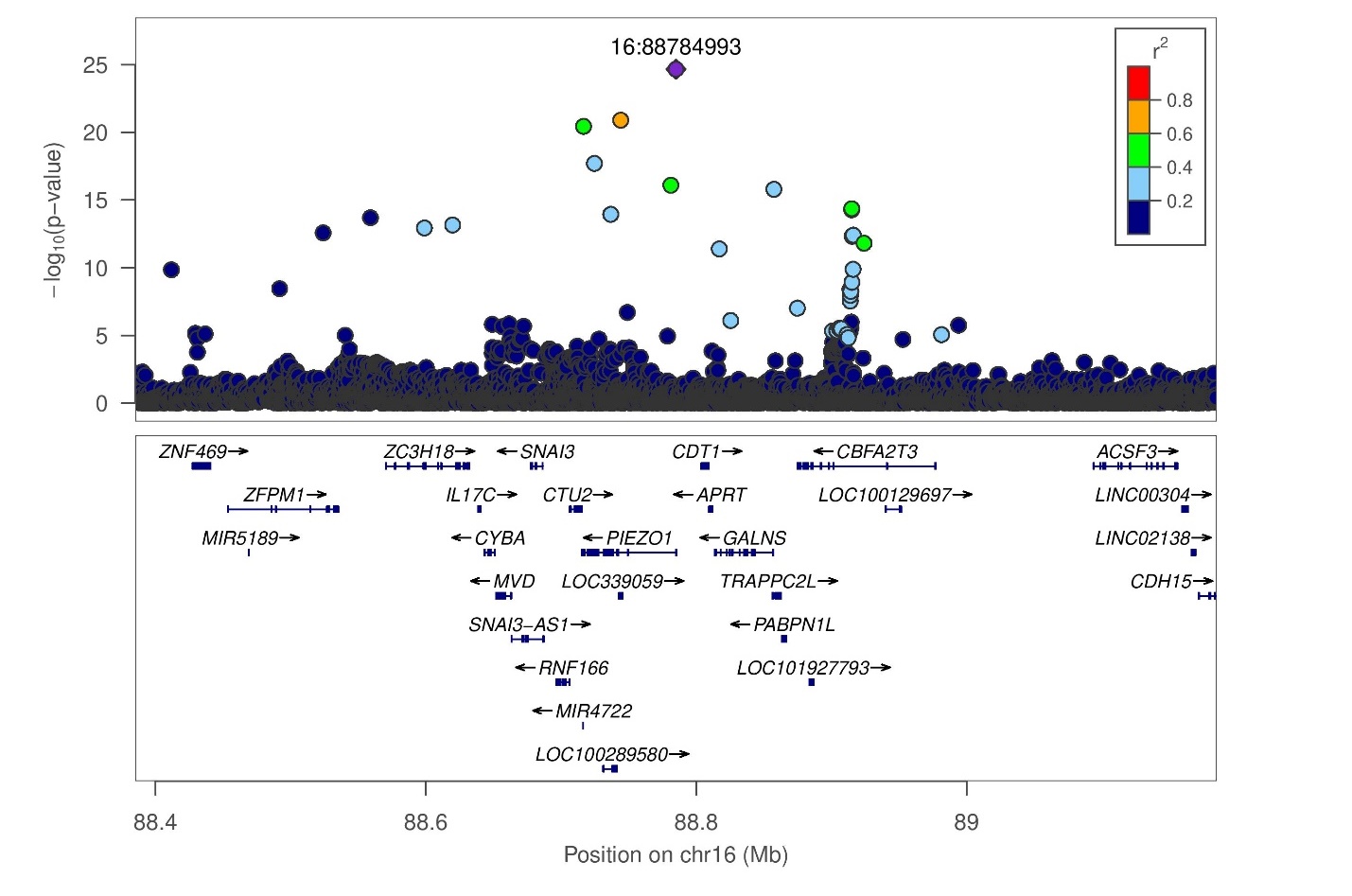

Note, our sentinel variant rs556126054 is the lead variant in unconditioned results. There is evidence that this signal is distinct from *PIEZ01* coding variant rs563555492. After conditioning on this lead variant, rs563555492 has p-value=0.002, but after conditioning on rs563555492, our sentinel rs556126054 still has a p-value=1e-7.

o. rs5030868, glycated hemoglobin, South Asian ancestry

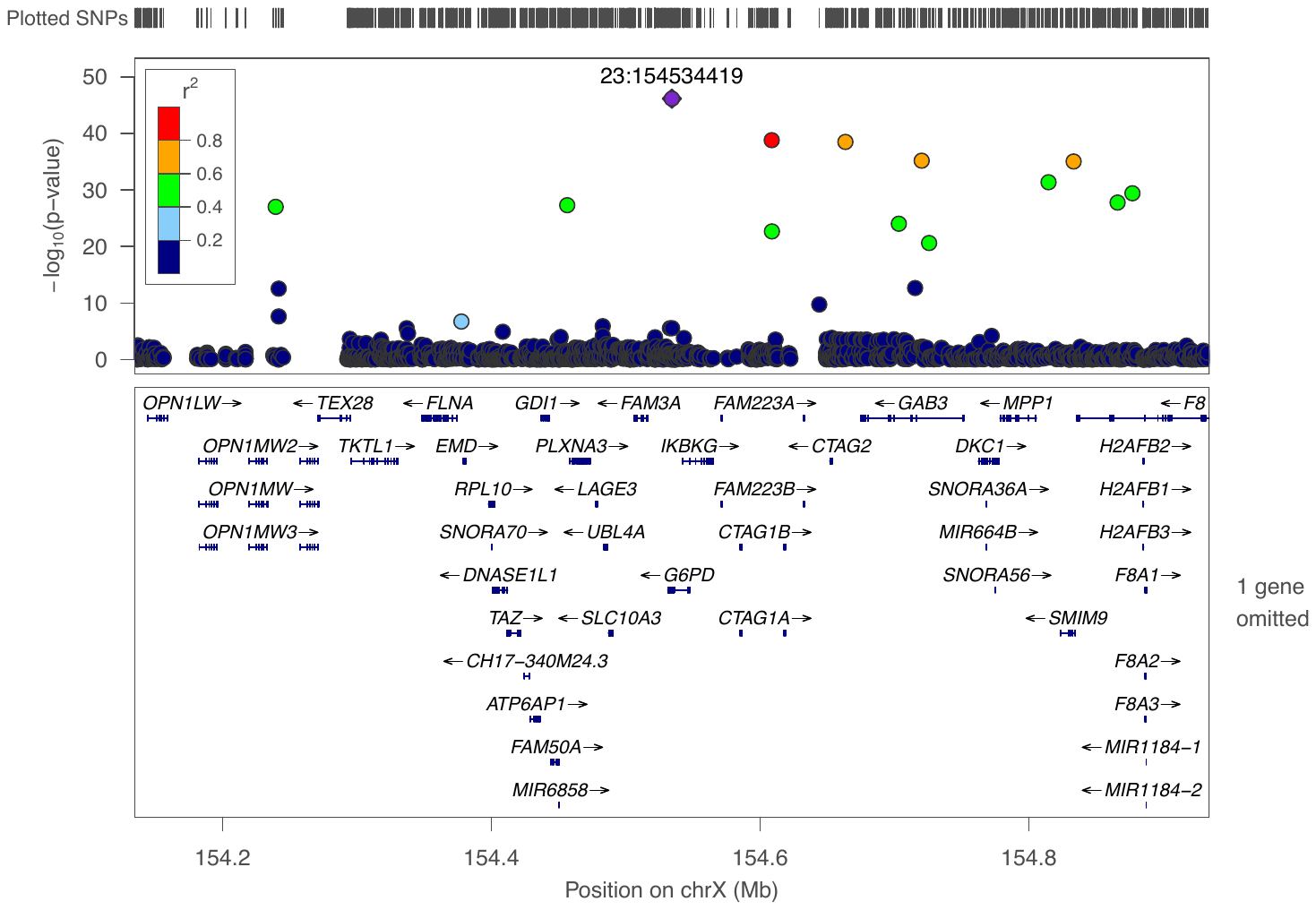

p. rs34680334, insulin-like growth factor 1, African ancestry

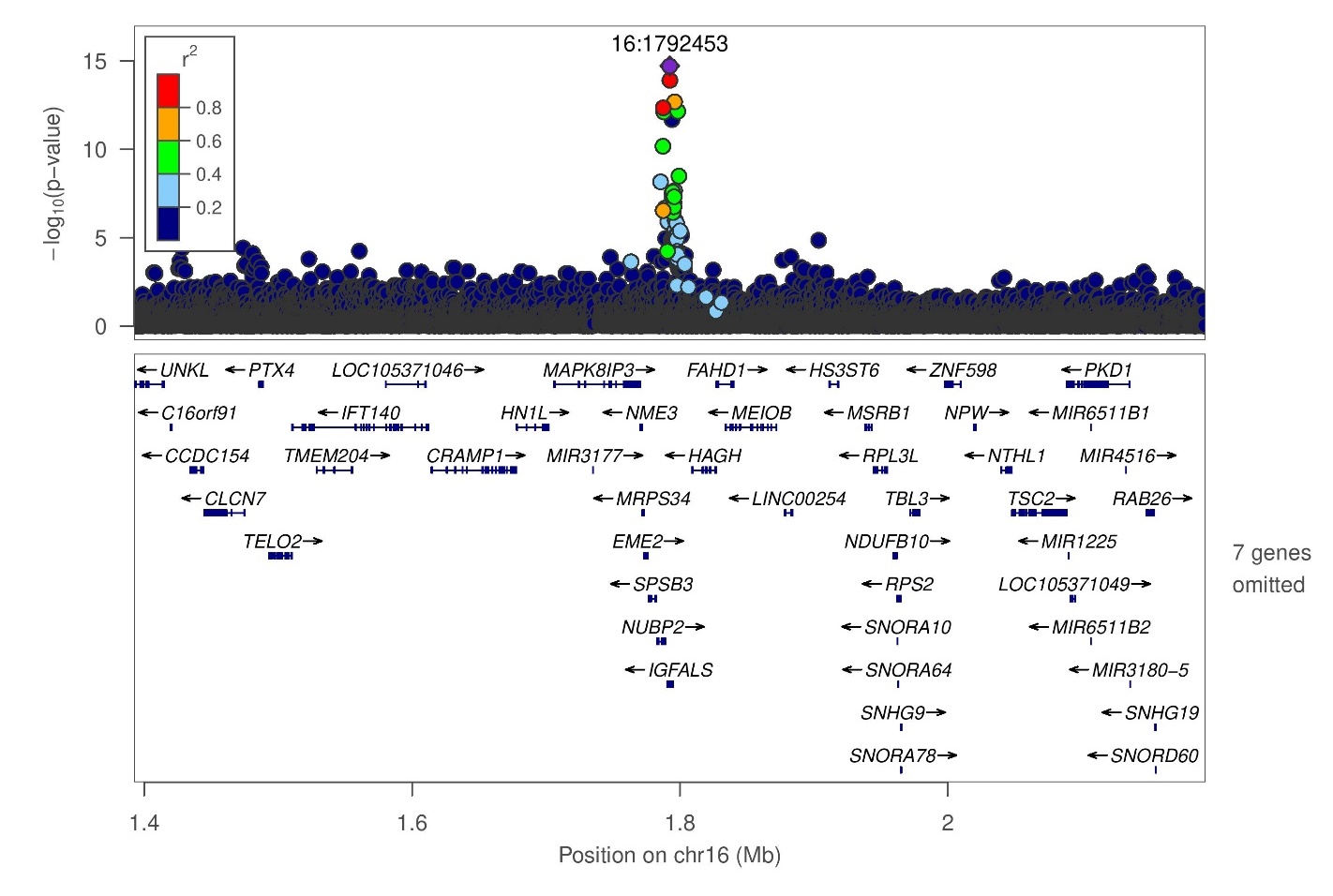

q. rs41270996, lipoprotein A, African ancestry

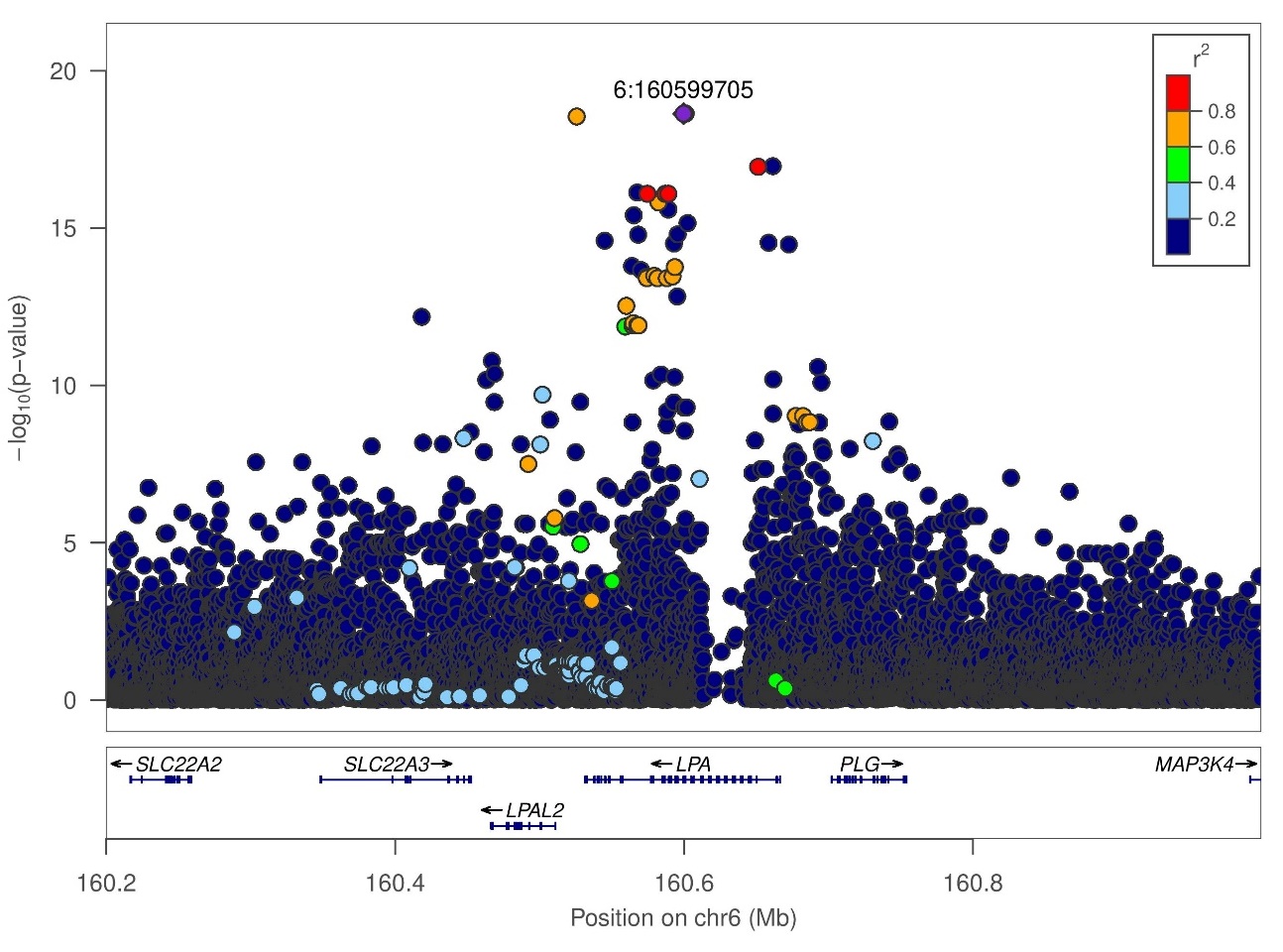

r. rs374112269, lipoprotein A, South Asian ancestry

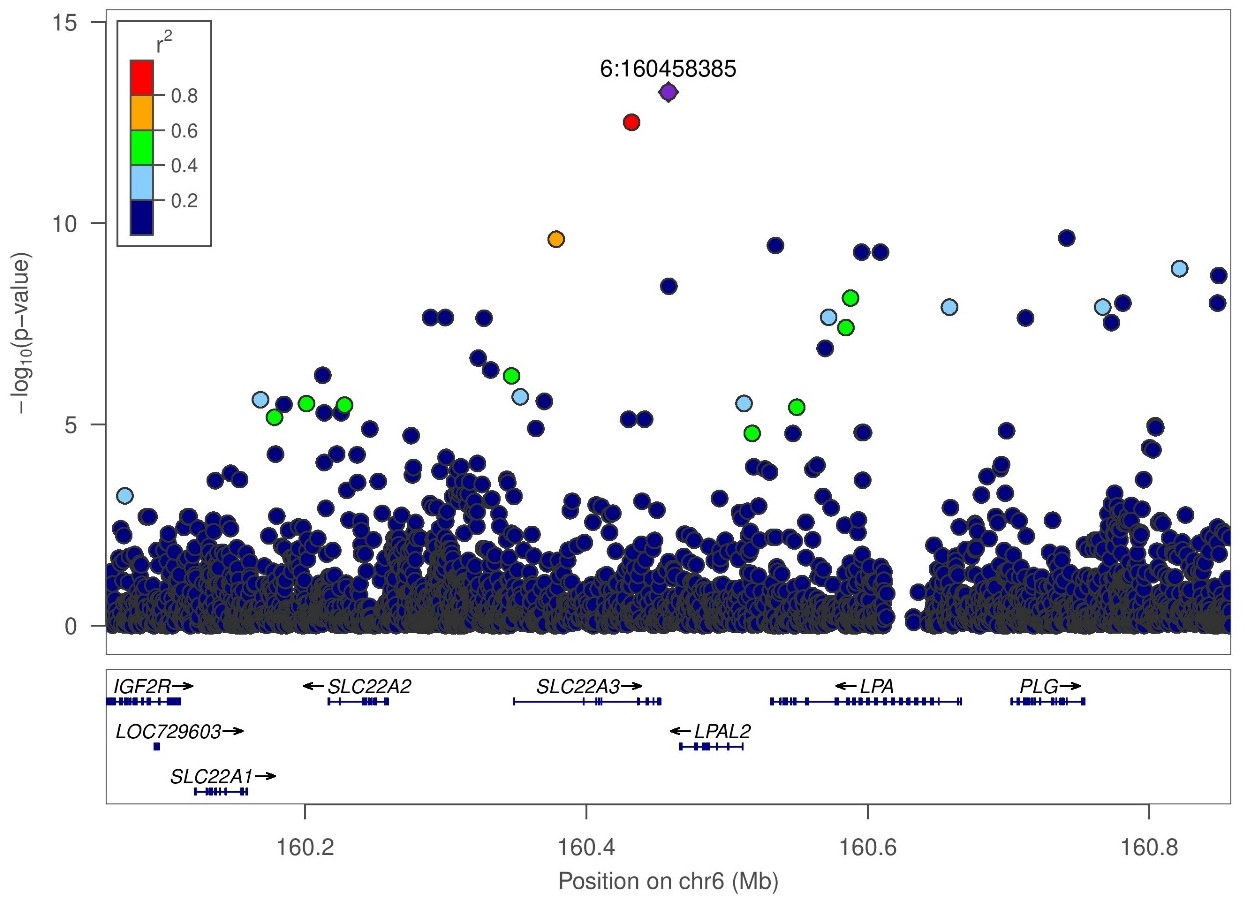

Figure S2: LocusZoom plots for conditionally distinct signals at loci with greater than one novel distinct signal.

a. Alkaline phosphatase African ancestry chromosome 6 locus, conditioned on all known GWAS variants from Table S2, Sinnot Armstrong et al. preprint (Table S3), and rs146351134

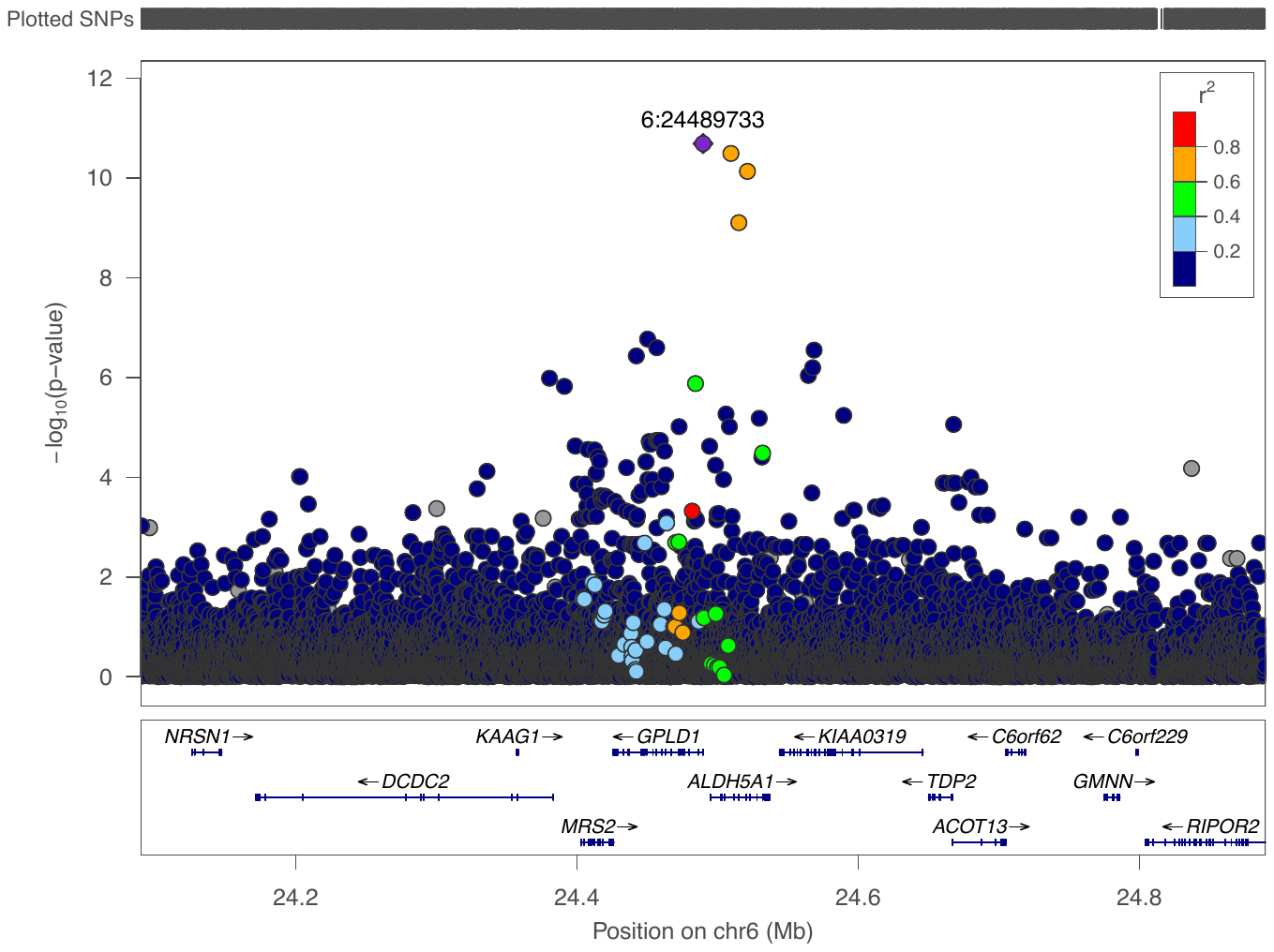

b. Lp(a) African ancestry chromosome 6 locus, conditioned on all known GWAS variants from Table S2, Sinnot Armstrong et al. preprint (Supplementary Table 3), and rs41270996

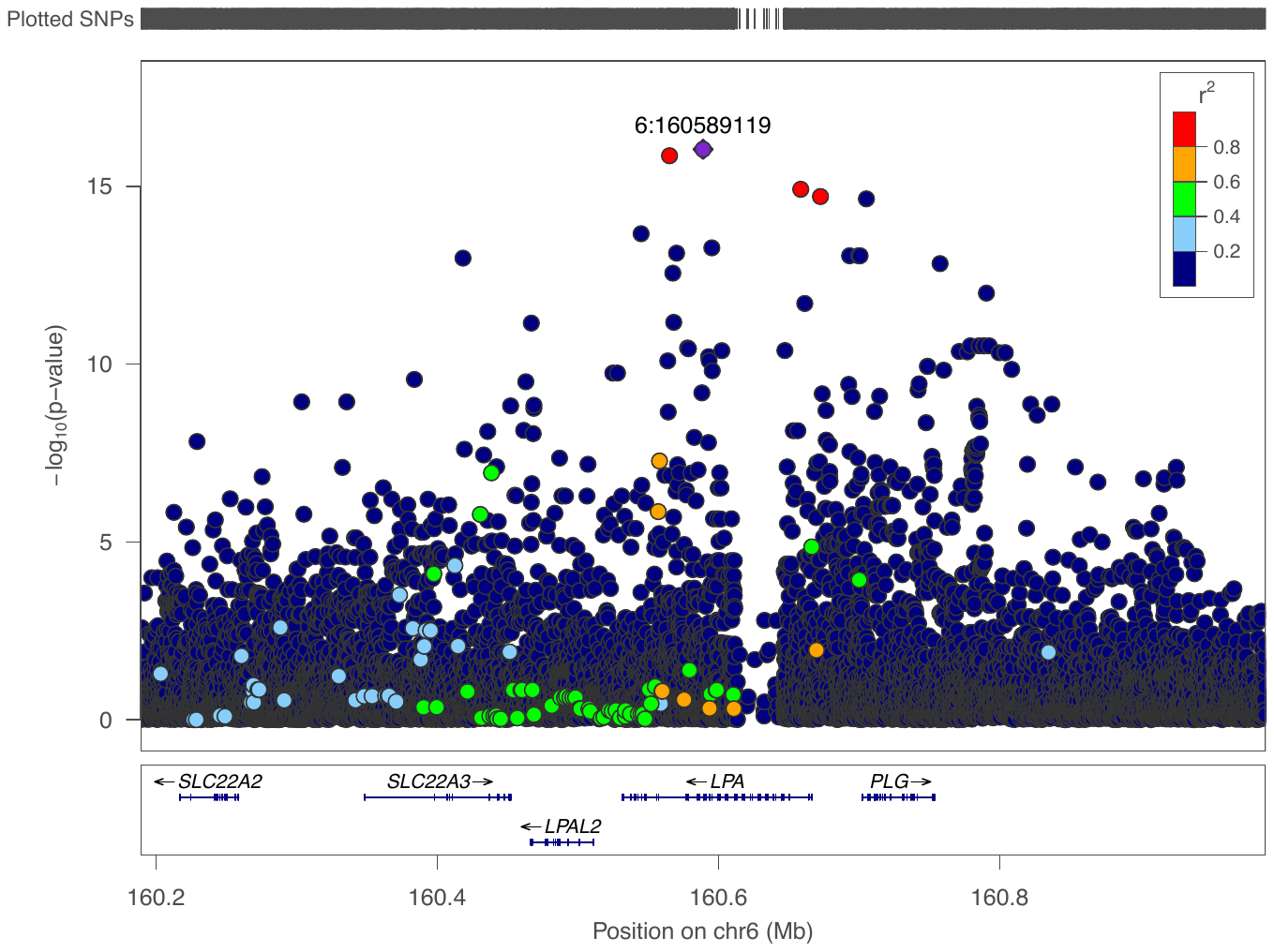

c. Lp(a) African ancestry chromosome 6 locus, conditioned on all known GWAS variants from Table S2, Sinnot Armstrong et al. preprint (Supplementary Table 3), and rs41270996, rs115739169.

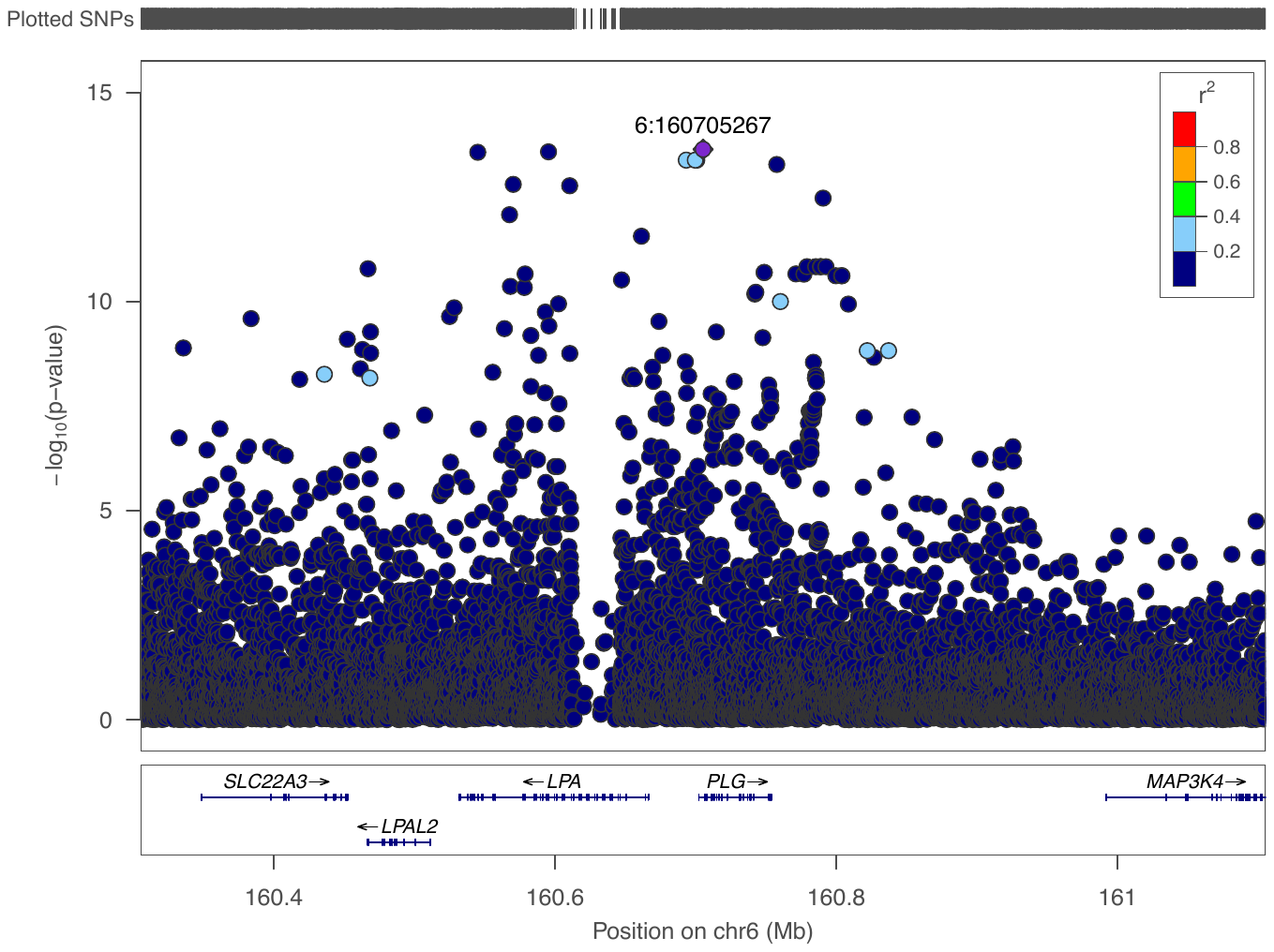

d. Lp(a) African ancestry chromosome 6 locus, conditioned on all known GWAS variants from Table S2, Sinnot Armstrong et al. preprint (Supplementary Table 3), and rs41270996, rs115739169 and rs186579824.

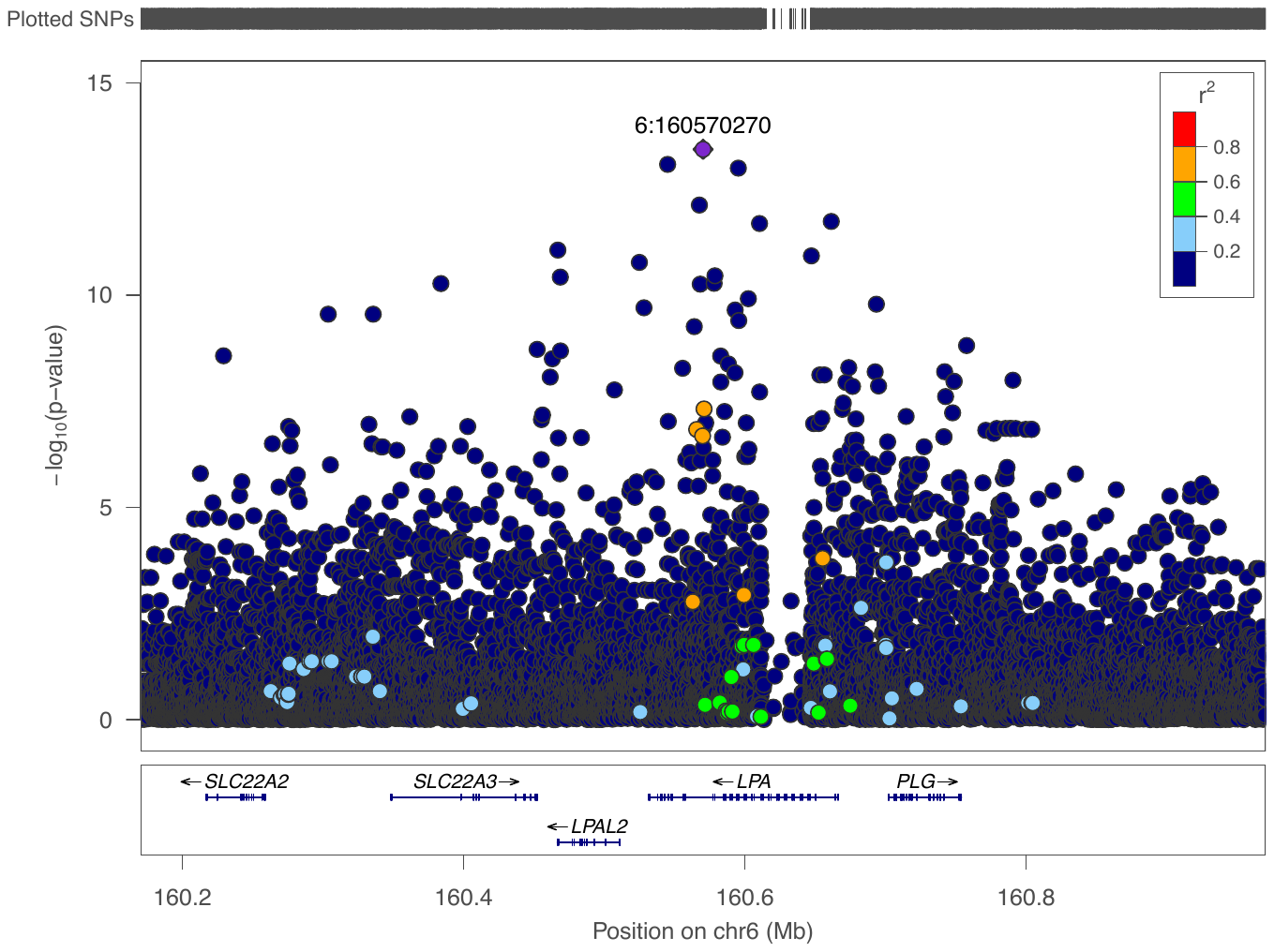

e. Lp(a) African ancestry chromosome 6 locus, conditioned on all known GWAS variants from Table S2, Sinnot Armstrong et al. preprint (Supplementary Table 3), and rs41270996, rs115739169, rs186579824 and rs138357319.

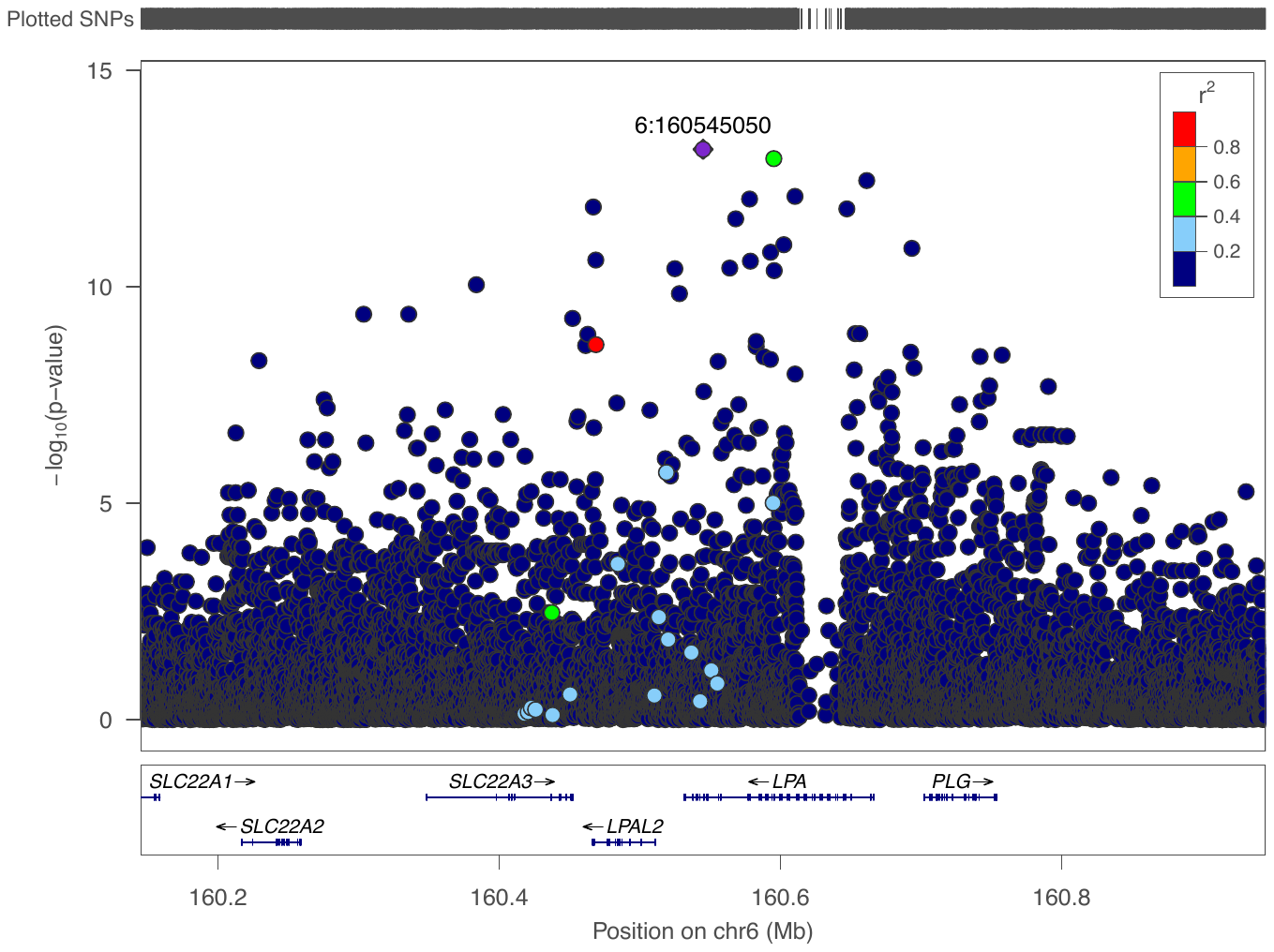

f. Lp(a) African ancestry chromosome 6 locus, conditioned on all known GWAS variants from Table S2, Sinnot Armstrong et al. preprint (Supplementary Table 3), and rs41270996, rs115739169, rs186579824, rs138357319 and rs150211198.

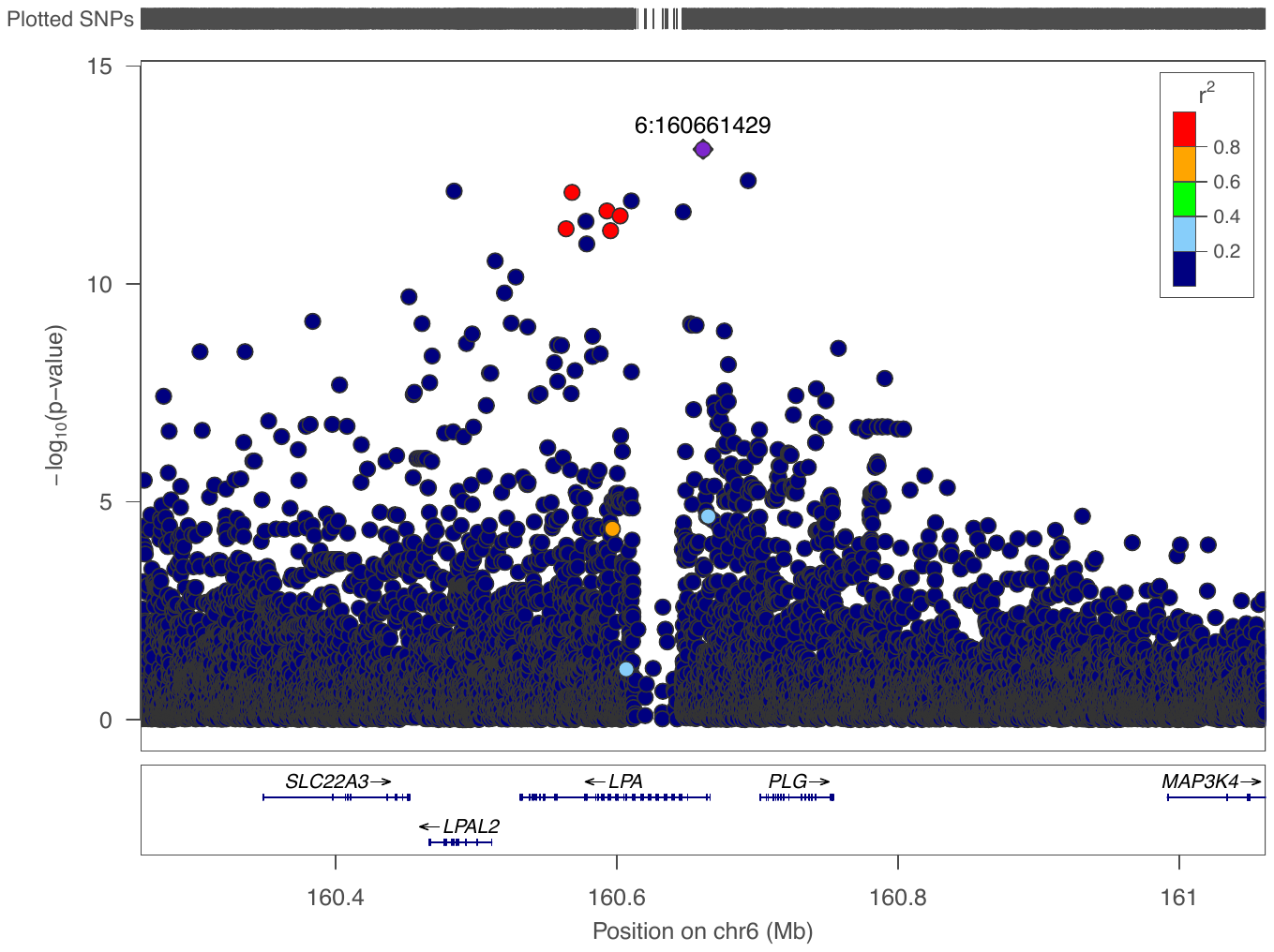

g. Lp(a) African ancestry chromosome 6 locus, conditioned on all known GWAS variants from Table S2, Sinnot Armstrong et al. preprint (Supplementary Table 3), and rs41270996, rs115739169, rs186579824, rs138357319, rs150211198 and rs1652507.

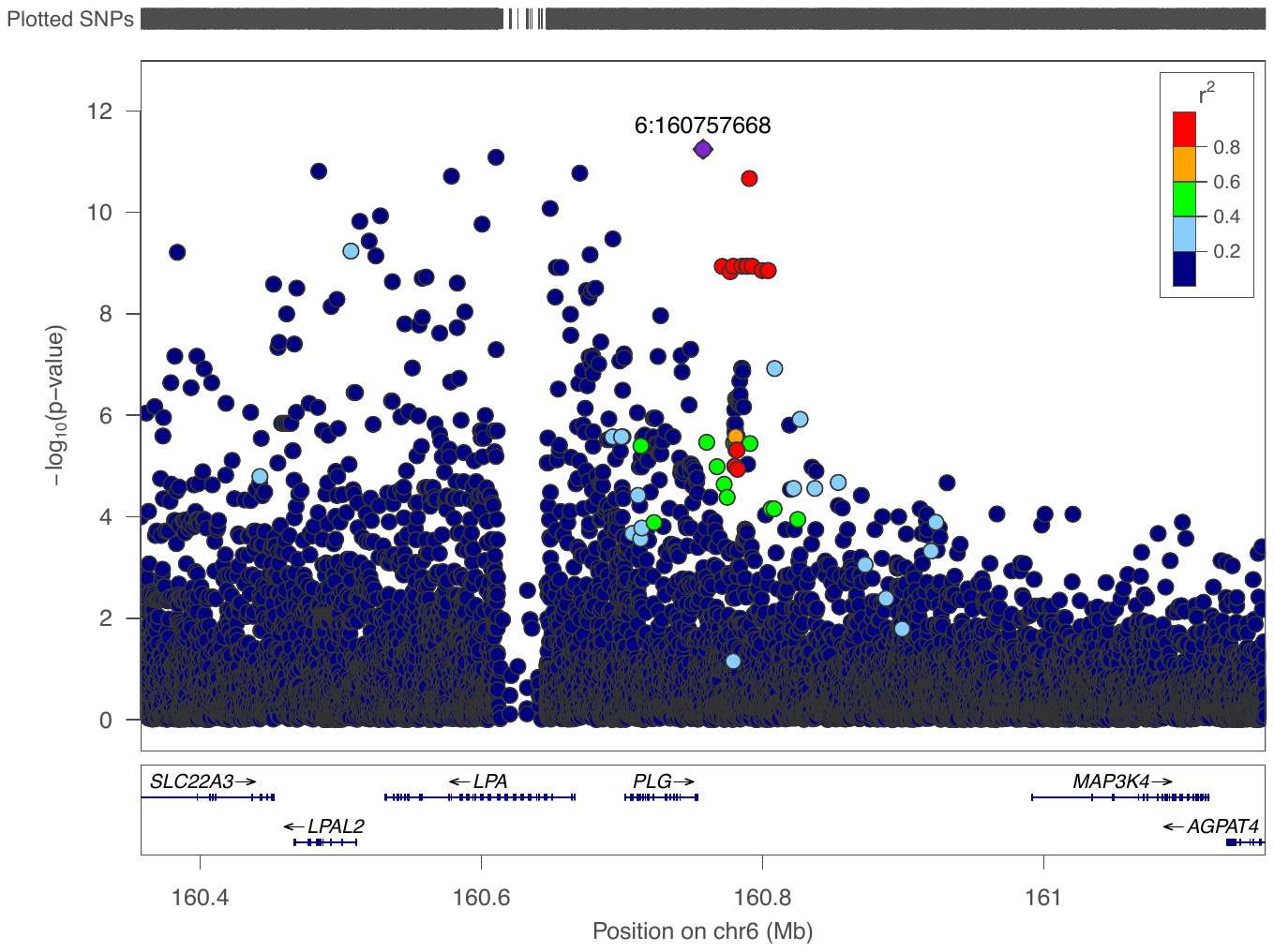

h. Lp(a) African ancestry chromosome 6 locus, conditioned on all known GWAS variants from Table S2, Sinnot Armstrong et al. preprint (Supplementary Table 3), and rs41270996, rs115739169, rs186579824, rs138357319, rs150211198, rs1652507 and rs60585169.

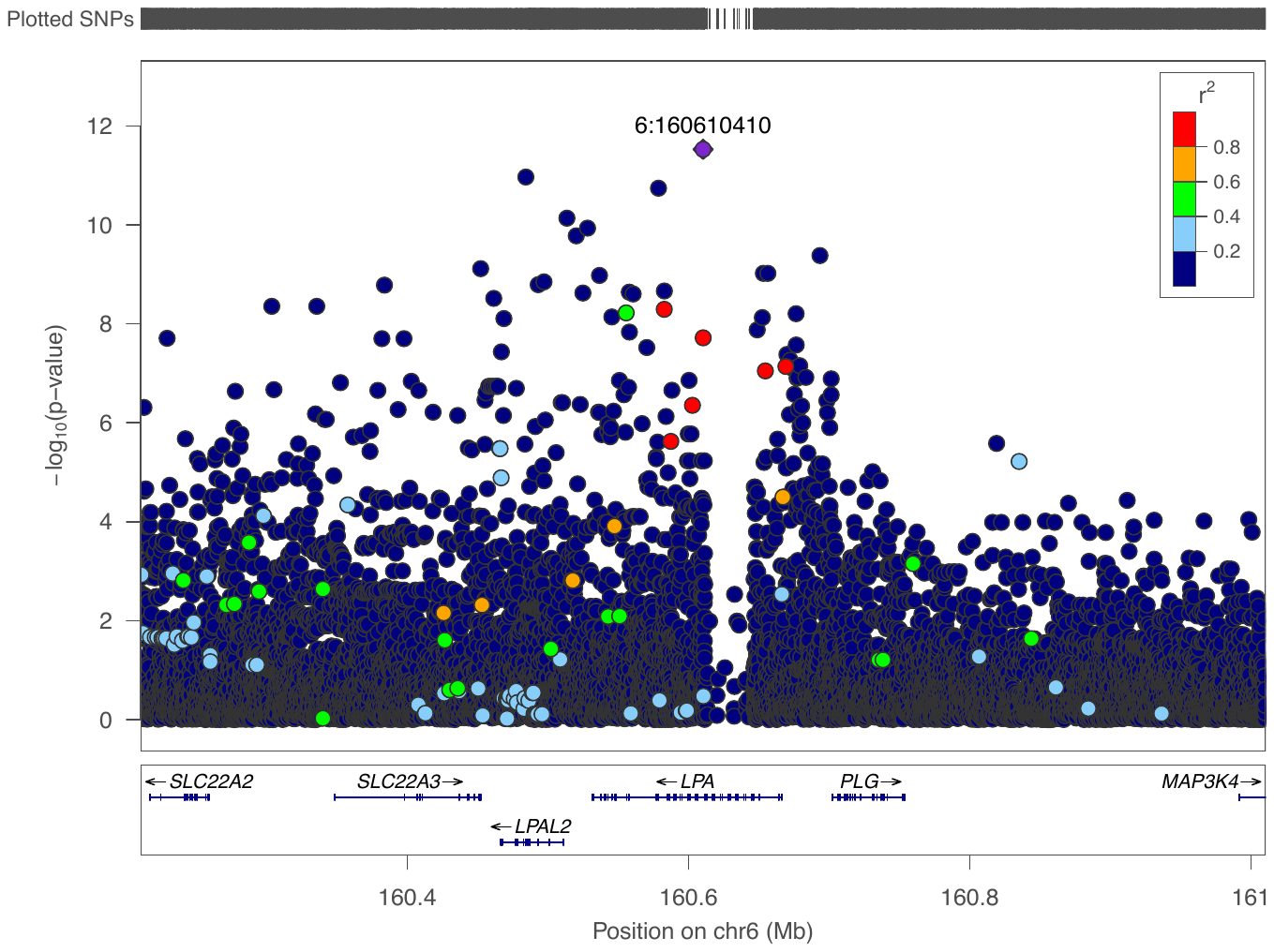

i. Lp(a) African ancestry chromosome 6 locus, conditioned on all known GWAS variants from Table S2, Sinnot Armstrong et al. preprint (Supplementary Table 3), and rs41270996, rs115739169, rs186579824, rs138357319, rs150211198, rs1652507, rs60585169 and rs141477332.

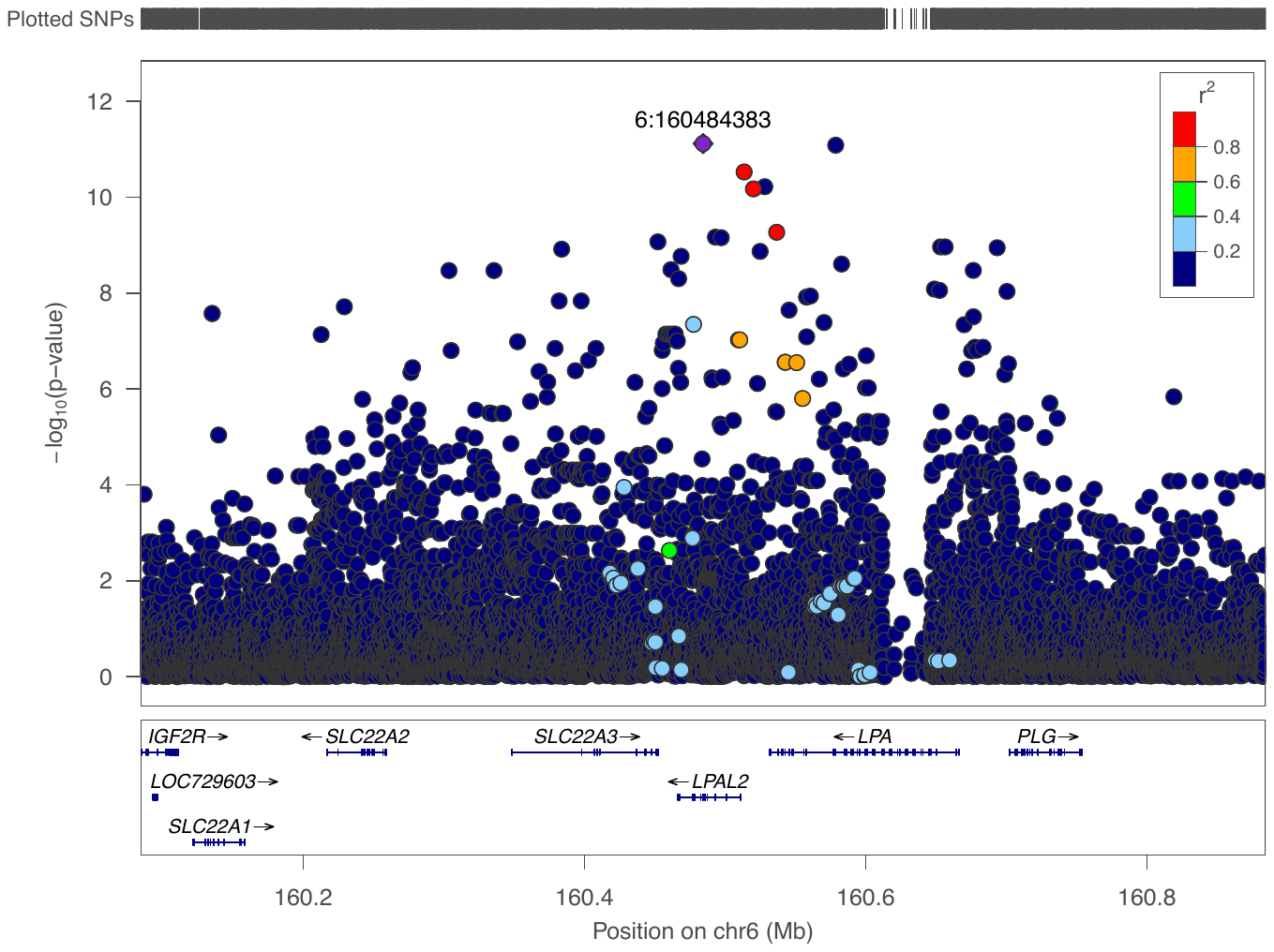

j. Lp(a) African ancestry chromosome 6 locus, conditioned on all known GWAS variants from Table S2, Sinnot Armstrong et al. preprint (Supplementary Table 3), and rs41270996, rs115739169, rs186579824, rs138357319, rs150211198, rs1652507, rs60585169, rs141477332 and rs1009324.

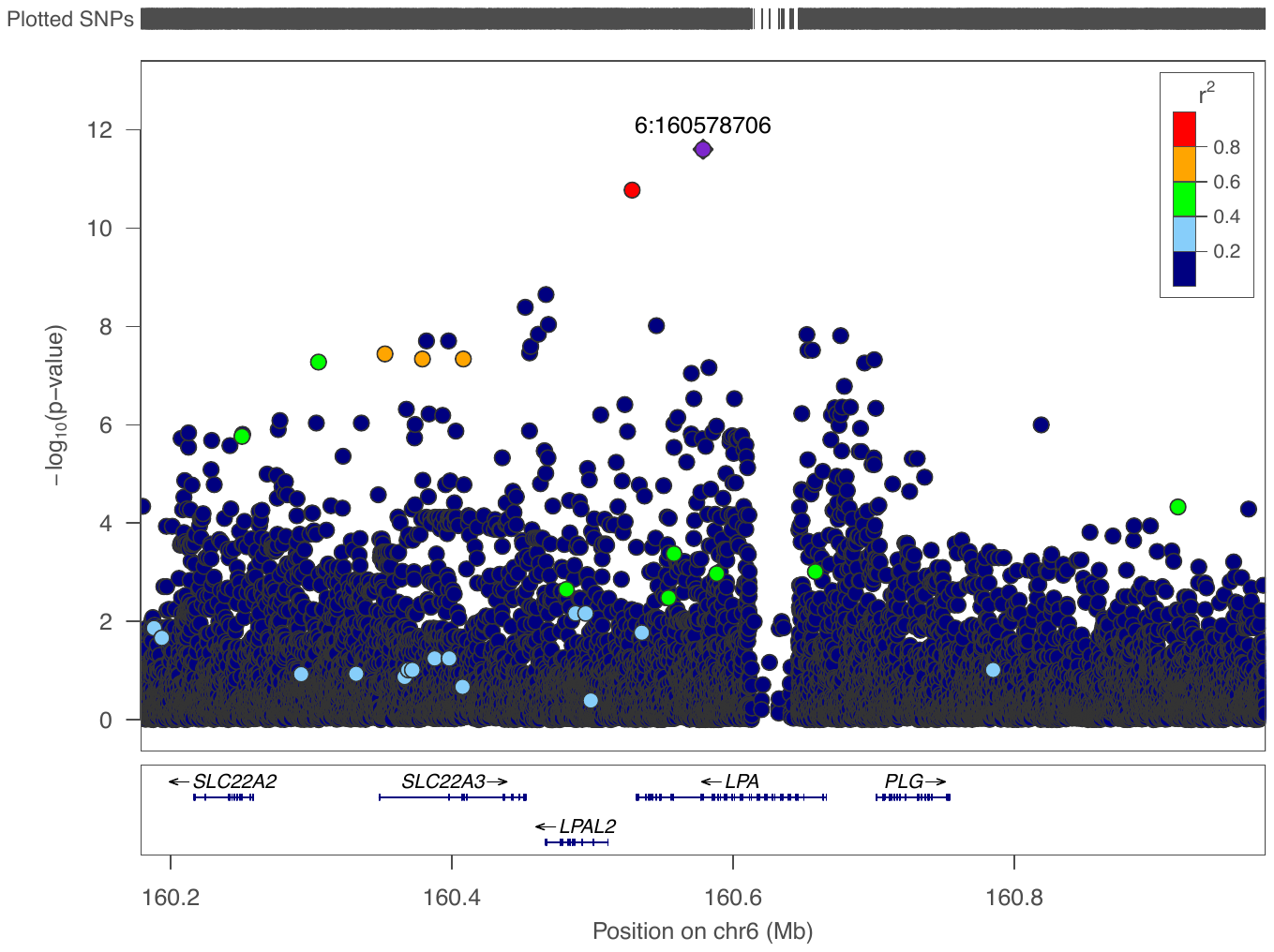

k. Lp(a) South Asian ancestry chromosome 6 locus, conditioned on all known GWAS variants from Table S2, Sinnot Armstrong et al. preprint (Supplementary Table 3), and rs374112269.

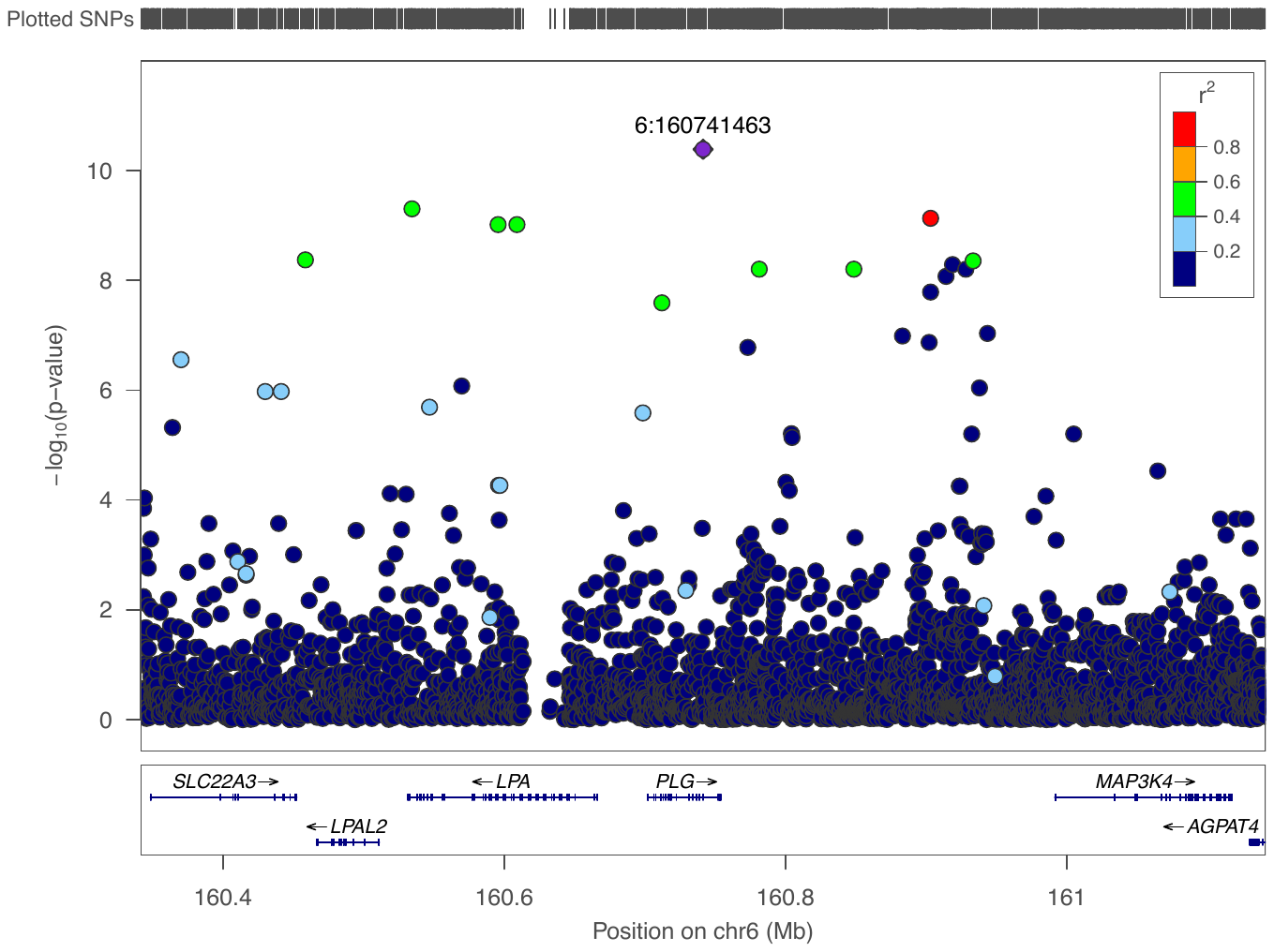
